# Circadian control of dopaminergic signaling to the mushroom body regulates sleep through rhythmic *Pka-C1* transcription in *Drosophila*

**DOI:** 10.1101/2025.08.29.673050

**Authors:** Blanca Lago Solis, Emi Nagoshi

**Affiliations:** Department of Genetics and Evolution and Institute of Genetics and Genomics of Geneva (iGE3), University of Geneva, CH-1205 Geneva, Switzerland

**Keywords:** dopamine, circuit, sleep, Drosophila, circadian rhythms, transcription, PKA-C1, mushroom body

## Abstract

Despite the progress in understanding the circadian pacemaker, the specific mechanism by which it regulates sleep remains incompletely understood. We have previously demonstrated that a substantial number of genes are rhythmically expressed in the mushroom body (MB) Kenyon cells (KCs), including *Pka-C1*, which encodes the catalytic subunit of protein kinase A (PKA). PKA-C1 plays a crucial role in promoting daytime wakefulness; however, the underlying mechanism remains elusive. Here, we show that the γ-lobe is the primary site of rhythmic *Pka-C1* expression using a newly developed *in vivo* luciferase reporter. Through a combination of *in silico* analysis, CRISPR mutagenesis, and chromatin immunoprecipitation, we identify the transcription factor Onecut as a regulator of *Pka-C1* transcriptional rhythms in γ-KCs. Furthermore, genetic trans-synaptic connectivity mapping and neuronal activity imaging reveal that the dorsal Lateral clock Neurons (LNds) provide inhibitory input to a subset of dopaminergic (DA) neurons in the protocerebral anterior medial (PAM) cluster, PAM-γ5, rhythmically modulating their activity. This, in turn, rhythmically activates MB γ-KCs via excitatory Dop1R signaling. Resulting γ-neuron activity rhythms drive *Pka-C1* transcriptional rhythms through Onecut. Furthermore, these PKA-C1 rhythms reinforce neuronal activity rhythms, creating a feedback cycle between transcriptional and neural activity rhythms that promote daytime wakefulness. Our findings highlight the conserved role of DA in promoting wakefulness and offer mechanistic insights into its complex regulation. More generally, this work provides a mechanistic framework for how circadian rhythms are translated into neural activity to orchestrate complex behaviors like sleep.

## INTRODUCTION

Circadian rhythms are biological processes that follow a cycle of approximately 24 h, governed by internal biological clocks. These clocks have been identified across a wide range of organisms, from cyanobacteria to humans, and regulate numerous daily behaviors and physiological processes, such as learning and memory formation, metabolism, locomotion, cognition, sleep, developmental timing, feeding, and courtship (Abbott & Zee, 2019; Campbell & Tobler, 1984; Cavanaugh et al., 2014; Jabbur et al., 2024; Lago Solis et al., 2025; Peschel & Helfrich-Förster, 2011; Woelfle et al., 2004). Circadian clocks enhance organisms’ fitness by enabling adaptation to daily environmental changes. Disruption in these rhythms is associated with an increased risk of neurological disorders such as depression, sleep disturbances, neurodegenerative diseases, and cognitive impairments(Abbott & Zee, 2019; Duret & Nagoshi, 2025; Zordan & Sandrelli, 2015). Understanding the interaction between circadian rhythms and brain function has therefore emerged as a critical area of research.

*Drosophila melanogaster*, exhibiting a diverse repertoire of complex behavior, coupled with its relatively simple nervous system and powerful genetic tools, provides an excellent model for exploring this connection. The *Drosophila* clock relies on the conserved interlocked transcriptional-translational feedback loops (TTFL). In the core of the TTFL the CLOCK/CYCLE (CLK/CYC) complex activates the expression of *period* (*per*) and *timeless* (*tim*) genes, and the PER-TIM protein complex accumulates in the cytoplasm during dusk, subsequently translocates to the nucleus to inhibit CLK/CYC transcriptional activity. A stabilizing loop involving PDP-1 and VRILLE (VRI) further regulates *Clk* transcription, contributing to the robustness of the circadian rhythm (Allada & Chung, 2009; Ashmore et al., 2003; Bargiello & Young, 1984; Glossop et al., 2003; Peschel & Helfrich-Förster, 2011).

In the adult fly brain, the molecular clock is present in approximately 150 neurons and 1800 glial cells, forming the central pacemaker. The clock neurons are organized into seven subclasses: three Dorsal Neuron clusters (DN1, DN2, and DN3) and five Lateral Neuron subclasses (LNd, s-LNv, 5th s-LNv, l-LNv, and LPN) (Muraro et al., 2013). Locomotor rhythms are regulated by the interaction of two primary oscillators: Morning (M) cells, composed of the s-LNvs (excluding the 5th s-LNvs), which release pigment-dispersing factor (PDF), and Evening (E) cells, including the LNds and the DN1s. M-cells drive morning anticipatory activity and free-running rhythms, while E-cells primarily govern evening activity peaks (Depetris-Chauvin et al., 2011; Muraro et al., 2013).

In addition to locomotor activity, flies display circadian rhythms in various behaviors, such as mating, courtship, learning, memory formation, feeding, and sleep-wake cycles (Barber et al., 2016; Chatterjee et al., 2010; Fropf et al., 2018; Kayser et al., 2014). The central clock neurons modulate other brain regions, including the mushroom body (MB), to rhythmically regulate these behaviors. The MB, the central information processing center in insects, plays a role in associative learning and state-dependent behaviors like sleep and hunger (Joiner et al., 2006; Modi et al., 2020). Each hemisphere of the MB comprises approximately 2000 Kenyon cells (KCs). Their dendritic arborizations form a calyx, and axons project together to form α/β, α′/β′, and γ-lobes. Those lobes can be further divided into 15 anatomical compartments (α1–3, β1–3, α′1–2, β′1–2, and γ1–5), distinguished by their gene expression profiles, neurotransmitter systems, and connectivity patterns (F. Li et al., 2020). This compartmentalization enables sensory information to influence behavior in a context-dependent manner.

The MB is densely innervated by modulatory neurons that release biogenic amines, such as dopamine, octopamine, and serotonin, inhibitory transmitters like GABA, and neuropeptides. These inputs, combined with cholinergic output from mushroom body output neurons (MBONs), form a circuit that shapes diverse behaviors (Aso, Sitaraman, et al., 2014; Driscoll et al., 2021; Mao & Davis, 2009). Dopaminergic (DA) neurons innervating the MB include those in the protocerebral posterior lateral (PPL1) and the protocerebral anterior medial (PAM) clusters, which play key roles in regulating behaviors, including learning (Kim et al., 2007), memory (Handler et al., 2019; Schwaerzel et al., 2003), and sleep (Driscoll et al., 2021; Kume et al., 2005; Mao & Davis, 2009). The interplay between DA neurons, MB compartments, and MBONs creates a dynamic network integrating sensory information to modulate behaviors. Given that behaviors controlled by the MB often follow circadian rhythms, the MB likely integrates time-of-day signals from clock neurons to coordinate its functions (Driscoll et al., 2021; Siju et al., 2021). However, the underlying mechanisms remain poorly understood.

We have previously demonstrated that a significant number of genes are expressed in a circadian manner in the MB, regulated by the molecular clock located in the central pacemaker neurons. These rhythmic genes include *Pka-C1*, which encodes the catalytic subunit of protein kinase A (PKA), and the tumor suppressor gene *Neurofibromin 1* (*Nf1*). Within the MB, NF1 activates PKA-C1 by increasing cAMP levels in a rhythmic fashion, which subsequently raises intracellular Ca^+2^ levels. The resulting activity rhythms in MB neurons, peaking during the day, promote daytime wakefulness (Machado Almeida et al., 2021). These findings underscore the extensive regulation of brain function by circadian clocks and their critical role in sleep regulation.

In this study, we investigate the molecular and neuronal mechanisms underpinning the control of the MB neurons by circadian pacemaker neurons. Using a newly developed *in vivo* luciferase reporter for *Pka-C1* gene expression, we identify the γ-KCs as the primary site of its rhythmic expression. Through a combination of RNAi, CRISPR mutagenesis, and chromatin immunoprecipitation (ChIP), we demonstrate that rhythmic *Pka-C1* expression is controlled by the transcription factor Onecut within the γ-KCs. Furthermore, genetic trans-synaptic connectivity mapping and neuronal activity imaging reveal that LNd clock neurons provide synaptic input to a subset of PAM DN neurons, specifically PAM-γ5 neurons projecting to the MB γ5 subdomain. LNds provide inhibitory input to PAM-γ5, while DA input from PAM-γ5 to γ-KCs is excitatory, mediated by dopamine D1-type receptor (Dop1R) signaling. Thus, rising LNd activity in the evening drives an elevation of calcium levels during the day and the middle of the night in PAM-γ5 neurons. This, in turn, triggers a similar temporal pattern of activity in γ-KCs, which drives the *Pka-C1* transcriptional rhythms. This circuit is further reinforced by the Pka-C1 rhythms feeding back to the γ-neuron activity rhythms. Our findings not only reveal a sleep regulatory circuit but also underscore the conserved role of DA in promoting wakefulness and its circadian control. Beyond the sleep regulatory network, this work expands our understanding of how temporal information is integrated into neural networks to shape daily patterns of behavior.

## RESULTS

### Tissue-specific *in vivo* luciferase reporter for *Pka-C1* expression

Rhythmic gene expression in the MB KCs can be regulated by transcriptional or post-transcriptional mechanisms. The latter includes modulation of mRNA stability via miRNA binding to the 3’ untranslated region (3’UTR) (Xue & Zhang, 2018; Yang et al., 2008). To investigate these possibilities, we focused on the *Pka-C1* gene, which exhibits clock-dependent circadian rhythms in expression in the MB neurons and plays a key role in mediating the MB’s regulation of sleep (Machado Almeida et al., 2021). We developed an *in vivo* luciferase reporter, comprising the 2107 bp *Pka-C1* promoter, an FRT-Stop-FRT cassette, the luciferase coding sequence, and a 3’UTR (Figure 1A). Two versions of the 3’UTR were utilized: the p10 3’UTR or the 3’UTR of the *Pka-C1* gene. This reporter system enables the recording of spatial and temporal activities of the *Pka-C1* promoter, in live, individual flies, by expressing the flippase (FLP) recombinase under a cell-type-specific GAL4 driver. Comparing two reporter versions (*Pka-C1-luc* and *Pka-C1-luc-3’UTR*) allows us to assess the contribution of the 3’UTR to these expression patterns.

**Figure 1.**
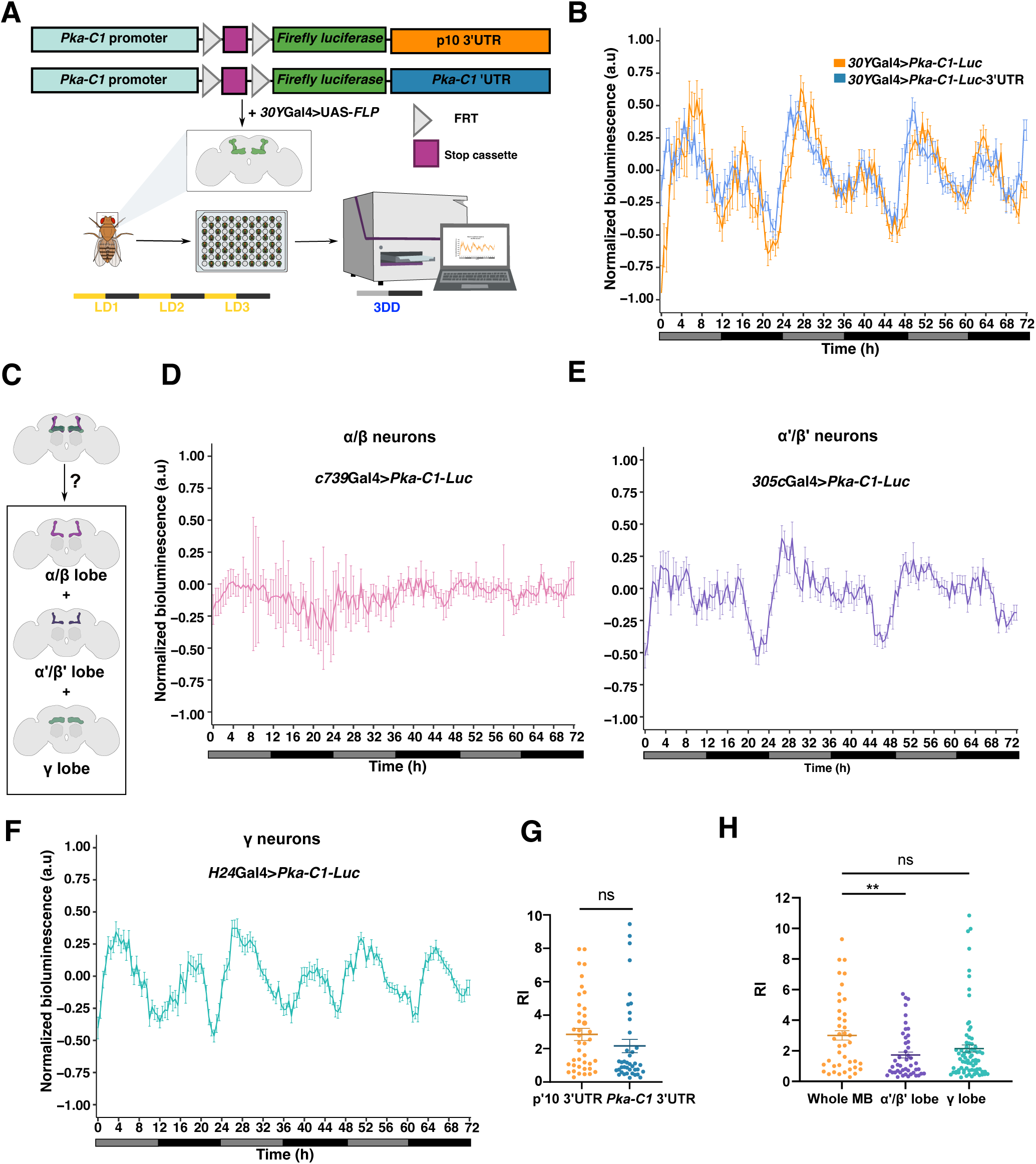
*Pka-C1* luciferase reporter reveals dominant transcriptional regulation *of Pka-C1* expression patterns in the MB γ-KCs. **A.** Tissue-specific in vivo luciferase reporters for *Pka-C1* expression and the bioluminescence recording system. Two versions of the reporter, one with a generic p10 3’UTR (*Pka-C1-luc*) and the other with the *Pka-C1* 3’UTR (*Pka-C1-luc*-3’UTR), were created. **B**. Luciferase expression patterns of the flies expressing the *Pka-C1-luc* reporter (orange) or *Pka-C1-luc*-3’UTR (blue) in the entire MB using *30Y*-GAL4. The graphs represent the median of the normalized bioluminescence ± SEM. n=55-60 flies per group. **C.** Schematic representation of the MB lobes. **D–F.** Bioluminescence signals from the *Pka-C1-luc* reporter in α/β-neurons (n=40) **(D**), α′/β′-neurons (n=85) (**E**), and γ-neurons (n=115) (**F**). The graphs represent the median of the normalized bioluminescence ± SEM. **G and H.** The rhythmicity index (RI) of *Pka-C1-luc* and *Pka-C1-luc*-3’UTR rhythms from the entire KC (**G**), and *Pka-C1-luc* rhythms in the entire KC populations, α′/β′-neurons, and γ-neurons (**H**). The center line and the error bars represent the mean ± SEM. Statistical comparisons were performed using a two-tailed t-test or Mann-Whitney test, depending on the data distribution. *p <0.05, **p<0.01, ***p<0.001, ****p<0.0001. ns, not significant.

As proof of principle, *UAS-FLP* was expressed throughout the entire MB in luciferase reporter flies using the pan-MB driver *30Y-GAL4* (Aso et al., 2009). The flies were entrained for three days under a 12-h light/12-h dark (LD) cycle and then placed individually in single wells of a 96-well plate containing food supplemented with luciferin (Figure 1A). Luciferase activity was recorded under constant darkness (DD) for the subsequent 74 h. Analysis of the bioluminescence data revealed that both reporters recapitulated the rhythmic *Pka-C1* gene expression profile (Machado Almeida et al., 2021), displaying a bimodal pattern (Figure 1B). Importantly, no significant differences in the percentage of rhythmic flies and bioluminescence profiles were observed between the two reporter lines (Table 1, Figure 1B and G). These results indicate that the promoter sequence introduced in the luciferase reporters is sufficient to drive the expression that mirrors the *Pka-C1* expression patterns. The 3’UTR sequence has little influence on these expression patterns, at least when analyzing summed bioluminescence across the entire MB KC populations.

### γ-neurons are the primary source of rhythmic *Pka-C1* expression in the MB

The MB consists of three distinct lobes, α/β, α′/β′, and γ, each exhibiting unique gene expression, connectivity, and behavioral roles. Given these anatomical and functional differences, *Pka-C1* expression patterns may vary across MB lobes (Figure 1C). To examine this potential difference, *Pka-C1*-*luc* reporter expression was driven in different subtypes of MB intrinsic KCs using the lobe-specific GAL4 drivers: *c739-GAL4* for the α/β-lobe neurons, *305c-GAL4* for the α′/β′-lobe neurons, and *H24-GAL4* for the γ-lobe neurons (Aso et al., 2009).

Strikingly, *Pka-C1*-*lu*c signals from the α/β-neurons lacked rhythmicity (Figure 1D), which was confirmed by the high percentage of arrhythmic flies (Table 1). This indicates that circadian modulation *Pka-C1* transcription does not occur in these cells. In contrast, luminescence from both the α′/β′- and γ-neurons displayed rhythms with two peaks within a 24-h day, similar to those observed in the entire MB KC population (Figures 1E and F). However, close comparison of the luminescence patterns in α′/β′-neurons and the entire KC population revealed a marked decrease in the second peak amplitude and significant reduction in the rhythmicity index (RI), an indicator of the strength of the rhythm, in α′/β′-neurons compared to the entire KCs (Figure 1H). Interestingly, the bioluminescence patterns from γ-neurons closely mirrored those of the entire MB (Figure 1F), showing no significant difference in RI (Figure 1H). These findings indicate that the γ-neurons are the primary contributors to the rhythmic expression of *Pka-C1* within the MB KC population.

While *Pka-C1* rhythms are primarily controlled by its promoter (Figure 1B), we wondered whether post-transcriptional modulation via the 3′UTR influences these rhythms in a lobe-specific manner. However, consistent with the observations of the entire KCs, luminescence patterns and RI between *Pka-C1-luc* and *Pka-C1-luc-3’UTR* reporters did not show significant differences in either α′/β′-(Supp. Figures 1A and B) or γ-neurons (Supp. Figures 1C and D). This suggests that post-transcriptional mechanisms via 3′UTR do not contribute to the modulation of *Pka-C1* gene expression in these neurons either.

In *per^0^* flies, bioluminescence rhythms from the *Pka-C1*-*luc* reporter in γ-neurons were abolished, confirming the dependence of *Pka-C1* rhythmicity on the circadian clock (Figure 2A and B, Table 1). Additionally, hyperactivation of γ-neurons with the thermosensitive TrpA1 channel at 29°C disrupted *Pka-C1* rhythms (Figures 2A, C-E, Table 1). Taken together with our previous findings that KCs exhibit clock- and PKA-C1-dependent activity rhythms (Machado Almeida et al., 2021), these results indicate that neuronal activity rhythms control circadian *Pka-C1* transcriptional rhythms in γ-neurons, and that Pka-C1 rhythms then feedback to reinforce activity rhythms.

**Figure 2.**
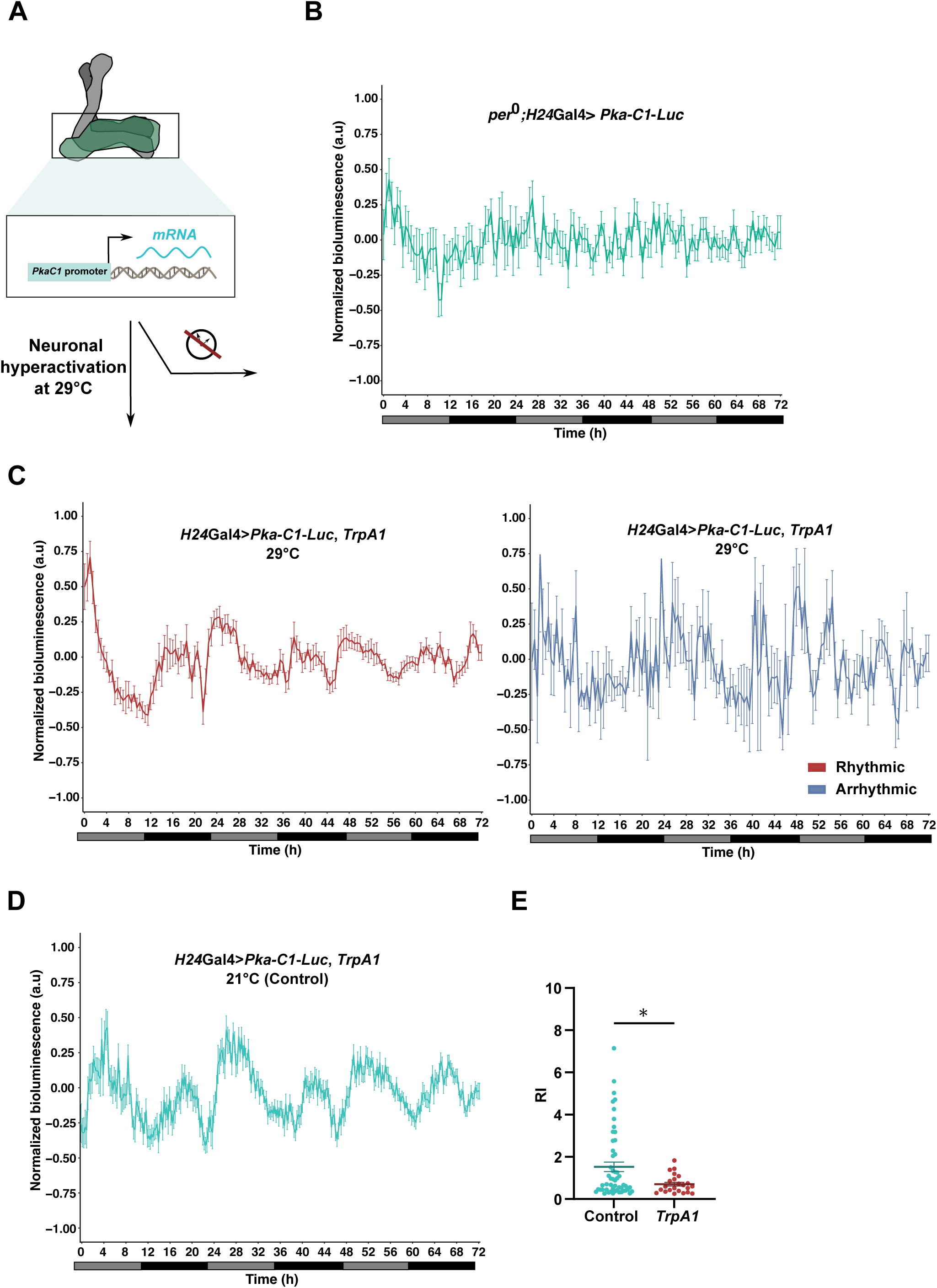
*Pka-C1* rhythms in γ-neurons depend on circadian clocks and neuronal activity dynamics. **A**. Schematic representation of the γ lobe and its contribution to the circadian modulation of *Pka-C1* gene expression. **B.** *Pka-C1-luc* reporter bioluminescence from γ-neurons in *per^0^* flies. n=26. (**C**). **C-E.** Hyperactivation of γ-neurons through the expression of *TrpA1* and temperature shift to 29°C impairs *Pka-C1-luc* rhythms. **C.** Bioluminescence patterns of *Pka-C1-luc* reporter of the rhythmic (red) and arrhythmic (blue) flies at 29°C. n=81. **D.** Bioluminescence patterns of *Pka-C1-luc* reporter at 21°C (control) flies. n=52. The middle line is the median, and the error bars represent the SEM. **E.** RI of the *Pka-C1-luc* signals from the control and TrpA1-hyperactivated flies. Mean ± SEM. Statistical comparisons were performed using t-test or Mann-Whitney test, depending on the data distribution. *p <0.05, **p<0.01, ***p<0.001, ****p<0.0001. ns, not significant.

### Transcription factor Onecut is required for *Pka-C1* transcriptional rhythms in γ-neurons

In our previous *in silico* analysis (Machado Almeida et al., 2021), a set of transcription factors (TFs) was identified as potentially involved in the circadian modulation of gene expression in the MB. Among these, four TFs – Onecut, Optix, Dorsal (dl), and Twist (twi) – have their predicted binding sites in the *Pka-C1* promoter (Figure 3A) and were shown to be expressed in the whole MB (Machado Almeida et al., 2021), including the γ lobe (H. Li et al., 2022). To test if these TFs are involved in the regulation of *Pka-C1* rhythmic expression, we first evaluated the efficiency of the available UAS-RNAi lines for each TF by quantitative PCR (qPCR) (Supp. Figure 2A). Although none of the tested RNAi lines achieved knockdown (KD) efficiency exceeding 50%, we selected the most effective RNAi for each gene for subsequent experiments. The *Pka-C1-luc* reporter and the RNAi line were then co-expressed in γ-neurons, and reporter bioluminescence was recorded.

**Figure 3.**
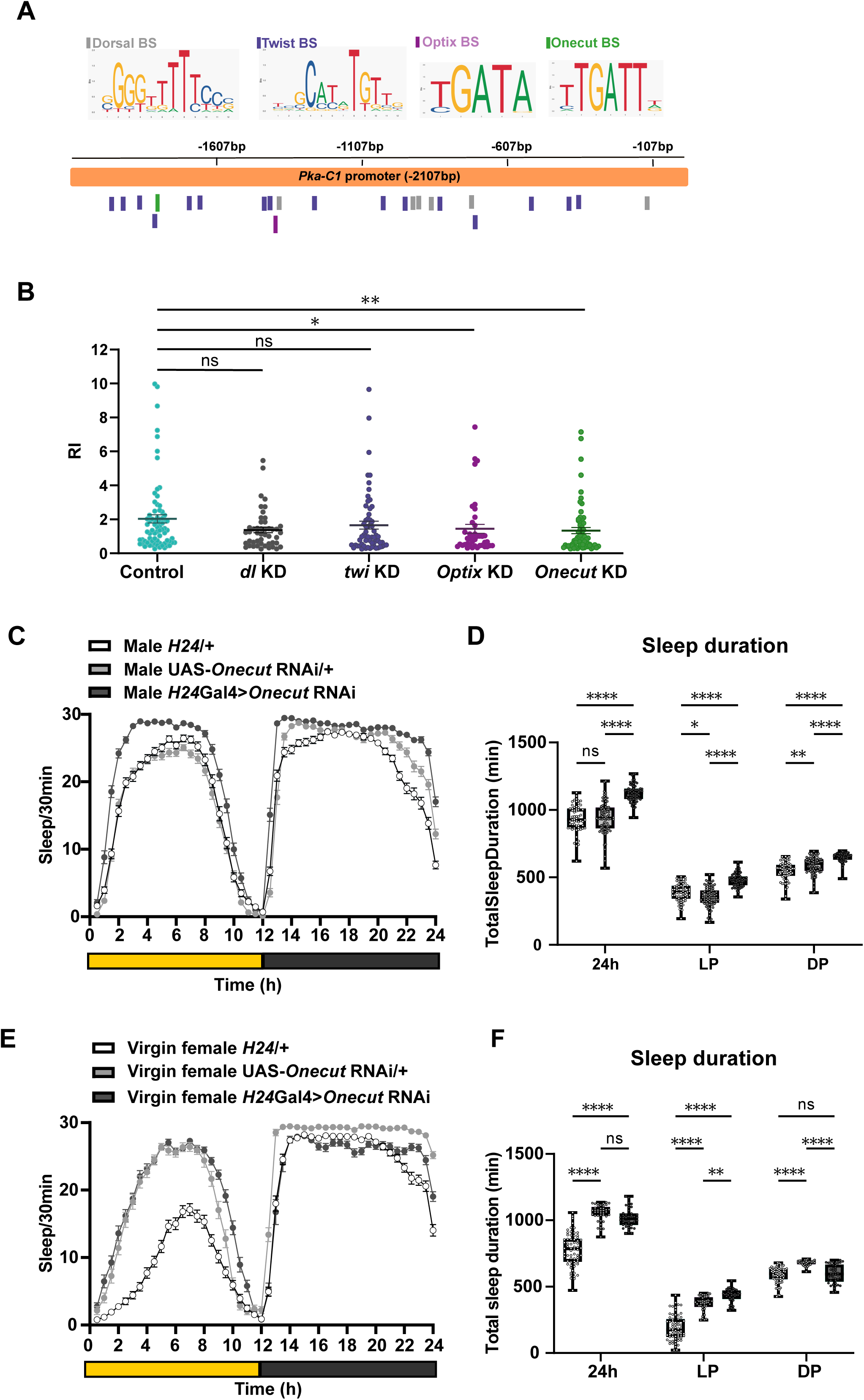
Onecut transcription factor regulates rhythmic expression of *Pka-C1* in γ – neurons and daytime wakefulness. **A.** Schematic representation of the different TF binding site positions in the *Pka-C1* promoter. **B.** The RI of the *Pka-C1-luc* bioluminescence in γ–KCs from control flies (*H24Gal4>pka-C1-luc*) and flies with KD of individual TFs. Mean ± SEM. Statistical comparisons were performed using a t-test or Mann-Whitney test, depending on the data distribution. n=63-105 per group. *p <0.05 and **p<0.01. ns, not significant. **C-F.** Sleep of the control flies carrying only the *H24* GAL4 driver and only the UAS-Onecut RNAi transgene, and the flies with *Onecut* RNAi driven with the same GAL4 were analyzed in males (n= 40-70 per genotype) (**C** and **D**) and virgin females (n= 30-40 per genotype) (**E** and **F**). *Onecut* KD increases sleep in both males and virgin females. Data are represented as mean values ± SEM in **C** and **E**. In **D** and **F**, the center lines of the boxplots indicate the median, the box boundaries are 25th and 75th percentiles, and the whiskers represent the minimum and maximum. LP, light period. DP, dark period. Two-way ANOVA with Dunnett’s correction for multiple comparisons was used for comparing genotypes. *p <0.05, **p<0.01, ***p<0.001, ****p<0.0001. ns, not significant.

RNAi of *dl* (Supp. Figure 2B and Figure 3B) and *twi* (Supp. Figure 2C and Figure 3B) did not result in significant changes in the bioluminescence patterns and RI, indicating that they are unlikely to be involved in the circadian transcriptional regulation of *Pka-C1*. However, RNAi of *Optix* (Supp. Figure 2D and Figure 3B) and *onecut* (Supp. Figure 2E and Figure 3B) reduced the RI, suggesting that these TFs play a role in controlling rhythmic expression of *Pka-C1*. Interestingly, Optix and Onecut have been reported to work in the same pathway in the auditory transduction mechanism in flies (Keder et al., 2020) and during the development of the retina in mice (Wu et al., 2013). This raises the possibility that they also act together in the circadian modulation of *Pka-C1* gene expression in the MB.

The single-cell RNAseq data in the Fly Cell Atlas (H. Li et al., 2022) indicate that *onecut* is expressed at significantly higher levels than *Optix* in γ-neurons. Moreover, our previous work demonstrated that *onecut* is rhythmically expressed in MB KCs under DD (Machado Almeida et al., 2021), consistent with its potential role as a regulator of rhythmic *Pka-C1* expression in the MB. Furthermore, we found that *onecut* KD in γ-neurons flies exhibited increased sleep in both males (Figure 3C and D) and virgin females (Figures 3E and F). This effect was more pronounced during the daytime, mirroring the phenotype observed in *Pka-C1* KD flies (Machado Almeida et al., 2021). These results strongly suggest that Onecut acts as a regulator of *Pka-C1* expression in γ-neurons, thereby contributing to the regulation of MB γ lobe function. Therefore, we focused on *onecut* in the subsequent study for dissecting the transcriptional regulation of *Pka-C1* expression.

Onecut is a transcription factor primarily known for its role in nervous system development in *Drosophila*, including neuronal differentiation and axon guidance. In adult *Drosophila*, Onecut continues to regulate genes essential for neuronal function and brain activity (Jauregui-Lozano et al., 2022; Nguyen et al., 2000). Given that *onecut* mRNA displays circadian oscillations and its potential involvement in the circadian regulation of *Pka-C1* gene expression, we hypothesized that its protein levels might also cycle across the day. To test this, we conducted an immunohistochemistry analysis of Onecut protein at six time points (ZT/CT0, 4, 8, 12, 16, and 20) in flies entrained under both LD and DD conditions to evaluate light dependency and circadian regulation, respectively (Figure 4A-C). Anti-Onecut fluorescence intensity was quantified in the cell bodies of γ-KCs, which were labeled with GFP using the γ-neuron-specific H24-Gal4 driver. The results revealed rhythmic Onecut protein levels in both LD and DD, with peaks at ZT4 and 16 in LD (Figure 4B), and CT4 and 12 in DD (Figure 4C). These peaks align with those of *Pka-C1*-*luc* rhythms (Figure 1B). In contrast, Onecut protein oscillation was completely abolished in *per^0^* in DD (Figure 4D). These findings indicate that Onecut cycling is under circadian clock control and consistent with the hypothesis that Onecut contributes to the circadian transcriptional regulation of *Pka-C1*.

**Figure 4.**
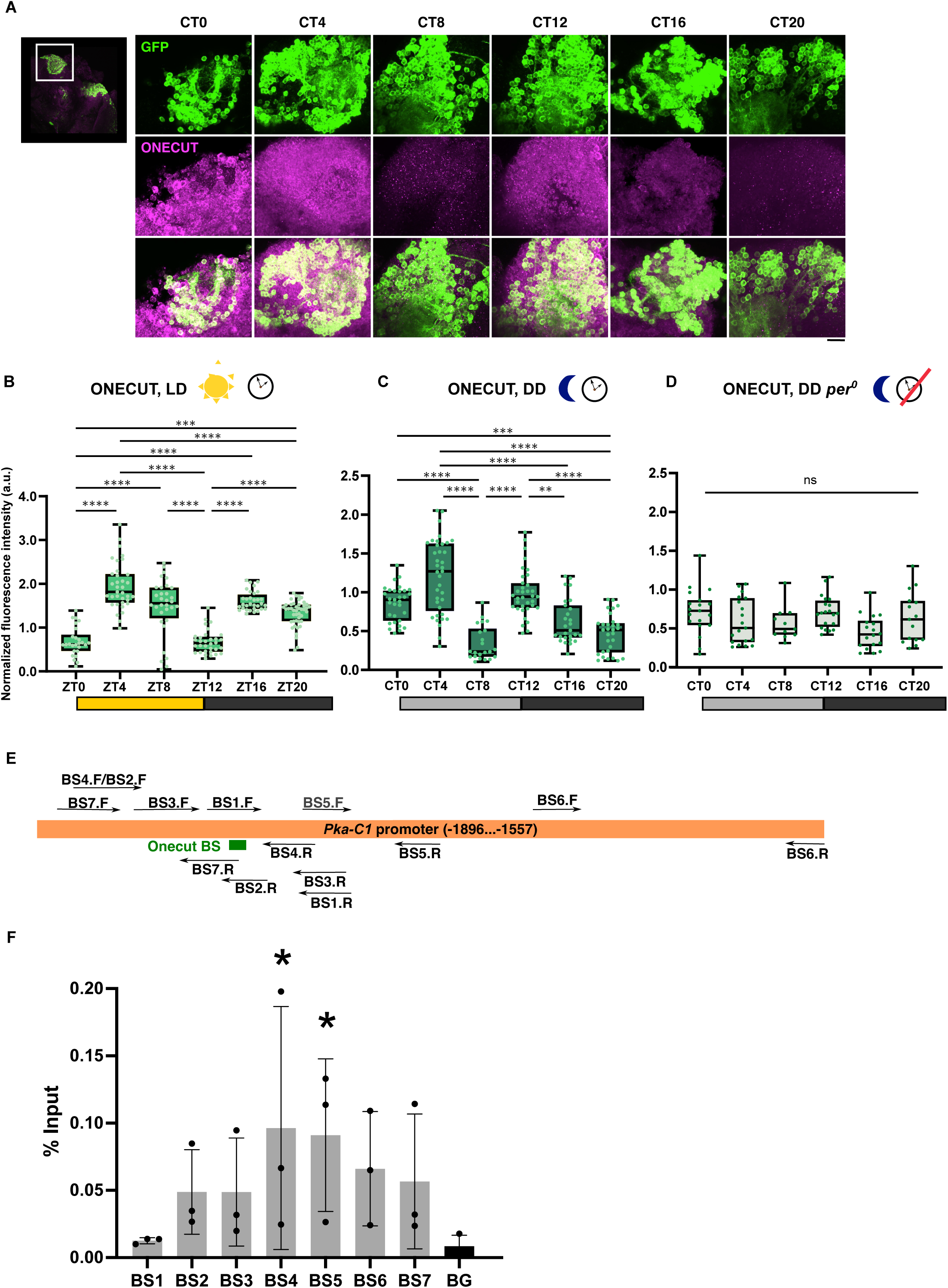
Onecut protein rhythmically accumulates in γ-neurons and binds the *Pka-C1* promoter. **A-C.** Immunostaining of fly brains with anti-Onecut and anti-GFP antibodies reveals clock- and light-dependent oscillations of Onecut accumulation in the soma of γ-neurons. GFP labels the cell bodies of γ-neurons using the *H24*-Gal4 driver. **A**. Representative confocal images of the brains of x40 objective with x4 zoom *H24>GFP* (*w^1118^* background) flies collected at 6 timepoints every 4 h in DD. Green, GFP. Magenta, Onecut. Scale bar, 10 μm. **B-D.** Quantification of the Onecut signals in the γ-neurons in control flies in LD (**B**) and DD (**C**) and in *per^0^* mutant flies in DD (**D**). The center lines of the boxplots indicate the median, and the whiskers represent the minimum and maximum. Dots represent individual values. arb. units, arbitrary units. n=30–40 per group. Based on the distribution of the data, the Kruskal–Wallis nonparametric test followed by Dunn’s multiple comparisons test, or the ordinary one-way ANOVA followed by Tukey’s multiple comparisons test was used for comparing values between timepoints. **E and F**. ChIP-qPCR analysis reveals Onecut binding to *Pka-C1* promoter. **E.** Schematic of the *Pka-C1* promoter in the regions −1896bp to −1557bp, the predicted Onecut core binding site, and the locations of qPCR primer pairs. **F**. ChIP-qPCR analysis of Onecut binding to the *Pka-C1* promoter. Enrichment is shown relative to input. Bar plots show the mean, error bars indicate the minimum and maximum values, and dots indicate individual replicates. Kruskal–Wallis nonparametric test followed by Dunn’s multiple comparisons test was used for comparing enriched regions. \**p* < 0.05, \*\**p* < 0.01, \*\*\**p* < 0.001, and \*\*\*\**p* < 0.0001. For clarity, only significant differences are indicated. Absence of a symbol denotes no significant difference.

Furthermore, we examined whether *Onecut* directly binds the *Pka-C1* promoter, using chromatin immunoprecipitation coupled to qPCR (ChIP-qPCR) with anti-Onecut antibodies. Genomic DNA from fly heads harvested at ZT4, when Onecut protein levels are highest (Figure 4B), was used as input DNA. Binding to or near the 7-bp predicted Onecut binding site within the *Pka-C1* promoter was analyzed by qPCR. This analysis revealed a significant increase in Onecut binding near the predicted binding site. Among the tested regions, the amplicons generated by the primer pairs BS4 and BS5 — located upstream and downstream of the 7-bp predicted motif, respectively — showed the highest enrichment. In contrast, the region directly covering the motif itself displayed lower enrichment (Figure 4E-F). These results indicate that Onecut preferentially associates with sequences flanking the predicted site, rather than the *in silico*-predicted binding motif itself. This may reflect binding to adjacent regulatory elements with higher affinity, or cooperative interactions with nearby cofactors.

### CRISPR mutants confirm Onecut’s role in regulating rhythmic *Pka-C1* expression and sleep

To further verify the role of Onecut in regulating *Pka-C1* expression and sleep, we sought to generate mutant flies with altered nucleotide sequences in or near the predicted Onecut binding site in the *Pka-C1* promoter. We used a germ cell-specific GAL4 driver to express the Cas9 enzyme and the gRNAs targeting the protospacer adjacent motifs (PAMs) near the Onecut binding site in the *Pka-C1* promoter to induce small insertions or deletions (indels) (Figure 5A). This strategy was applied to the fly strain heterozygous for the *Pka-C1-luc* reporter to generate mutations either in the endogenous *Pka-C1* gene promoter or in the promoter of the *Pka-C1-luc* transgene. This strategy yielded two different types of mutant lines: one carrying the wild-type *Pka-C1-luc* reporter but with a mutation in the endogenous *Pka-C1* promoter (mutant 67.4) and the other having a *Pka-C1-luc* reporter with a mutated promoter but no mutation in the endogenous *Pka-C1* (mutant 69.4) (Supp. Figures 3A-C). Importantly, both 67.4 and 69.4 mutants harbor indels within 12bp upstream of the predicted Onecut binding motif. Since the upstream flanking sequences showed the highest Onecut binding in our ChIP assay (Figures 4E and F), these mutants likely have reduced Onecut binding and, consequently, altered transcriptional regulation.

**Figure 5.**
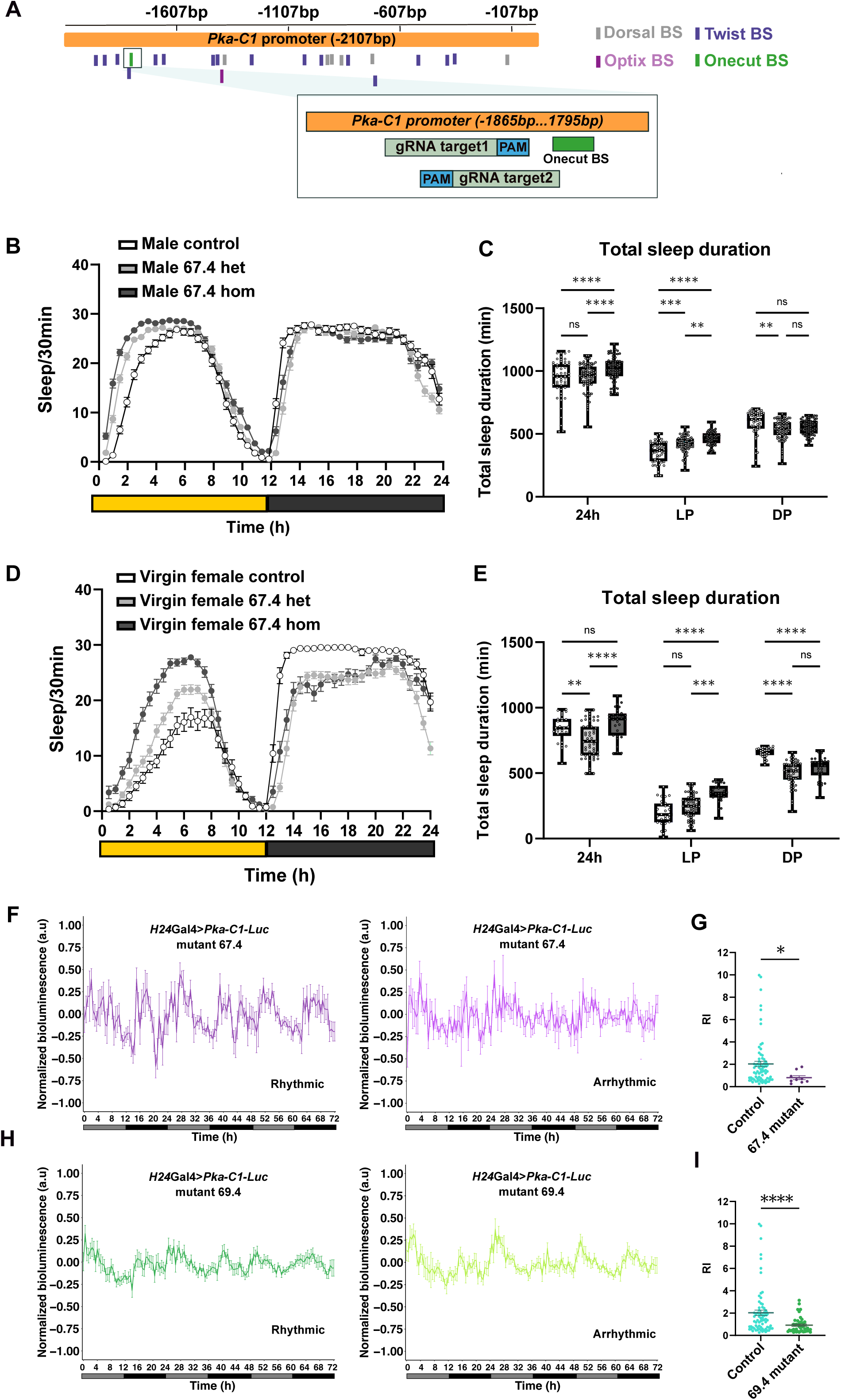
Onecut binding site mutations in the *Pka-C1* promoter increase sleep and disrupt *Pka-C1* expression rhythms in both *cis* and *trans*. **A.** Schematic representation of the different TF binding sites positions in the *Pka-C1* promoter and the CRISPR strategy targeting the Onecut binding site **B-E**. Sleep in males (**B and C**) and virgin females (**D and E**) of the 67.4 mutant line and controls. n=20-40 flies per genotype. In **C** and **E**, the center lines of the boxplots indicate the median, box boundaries are the 25th and 75th percentiles, and the whiskers represent the minimum and maximum values. Control, *w^1118^*. het, heterozygous mutant. hom, homozygous mutant. LP, light period. DP, dark period. The sleep data was analyzed with Two-way ANOVA with Dunnett’s correction for multiple comparisons. **F-I**. Bioluminescence recordings from the two CRISPR mutant lines. Line 67.4 (n=20) (**F** and **G).** Line 69.4 (n= 79) (**H** and **I**). Average bioluminescence plots (median ± SEM) of rhythmic and arrhythmic flies from each line are shown in the left and right panels, respectively (**F** and **H**). RI of rhythmic individuals from each mutant is compared with that of *H24*Gal4>*Pka-C1-luc* control flies (n=115). Mean ± SEM, Statistical comparisons were performed using a t-test or Mann-Whitney test, depending on the data distribution (**G** and **I**). *p <0.05, **p<0.01, ***p<0.001, ****p<0.0001. ns, not significant.

Sleep analyses revealed an increase in the total sleep duration in both males and females of the homozygous 67.4 mutant line compared to the heterozygotes and controls. The increase was specifically observed during the daytime (Figures 5B-E). Furthermore, bioluminescence recordings of the *Pka-C1*-*luc* reporter signals showed a significant decrease in rhythmicity in both mutants. Line 67.4, which carries a wild-type *Pka-C1-luc* reporter and homozygous mutations in the endogenous *Pka-C1* promoter, showed a significant decrease in both the percentage of rhythmic flies and RI (Figures 5F and G and Table 1). Likewise, line 69.4, which has a mutated *Pka-C1-luc* reporter but a wild-type *Pka-C1* gene, also displayed a significant reduction in rhythmic flies and RI (Figures 5H and I, Table 1). These results indicate that mutations in the *Pka-C1* promoter not only directly disrupt its gene expression rhythms in *cis* but also trans-dysregulate *Pka-C1* gene expression from a wild-type promoter. This *trans* effect is likely due to altered PKA-C1 rhythms, which disrupt neuronal activity rhythms (Machado Almeida et al., 2021). These findings collectively provide compelling evidence of a functional link between Onecut and *Pka-C1* within the MB γ-neurons.

### LNd clock neurons input time-of-day information to the MB γ lobe via PAM DA neurons

Our results so far indicate that *Pka-C1* expression in γ-neurons is rhythmically regulated by a transcriptional mechanism involving Onecut, likely modulated by neuronal activity dynamics. These dynamics are probably driven by time-of-day signals from clock neurons to the MB γ lobe. Based on the adult fly brain connectome (F. Li et al., 2020), a direct synaptic connection between clock neurons and the γ lobe seems improbable. However, the γ lobe is densely innervated by DA neurons from the PAM and PPL1 clusters (Aso, Hattori, et al., 2014). Additionally, our previous study demonstrated that subsets of PAM neurons receive direct synaptic input from clock neurons (Majcin Dorcikova et al., 2023). Therefore, PAM neurons are the likely intermediaries relaying circadian signals from clock neurons to the γ lobe.

Different PAM and PPL1 subpopulations innervate anatomically distinct γ lobe compartments, γ1 to γ5 (Figure 6A) (Aso, Hattori, et al., 2014). To determine connectivity between these DA neurons innervating the γ lobe and clock neurons, we used *retro*-Tango, a transsynaptic retrograde circuit mapping tool (Sorkaç et al., 2023). This system utilizes a ligand-receptor interaction at the synapse that triggers a signaling cascade, resulting in the expression of mtdTomato red fluorescent protein (RFP) in presynaptic and green fluorescent protein (GFP) in postsynaptic neurons. We targeted PAM or PPL1 subpopulations innervating different γ lobe compartments using specific split-GAL4 drivers (Figure 6B). Additionally, the γ-neuron driver was included to examine the possibility of a direct connection between clock neurons and γ-neurons. Clock neurons were identified using anti-PER immunostaining.

**Figure 6.**
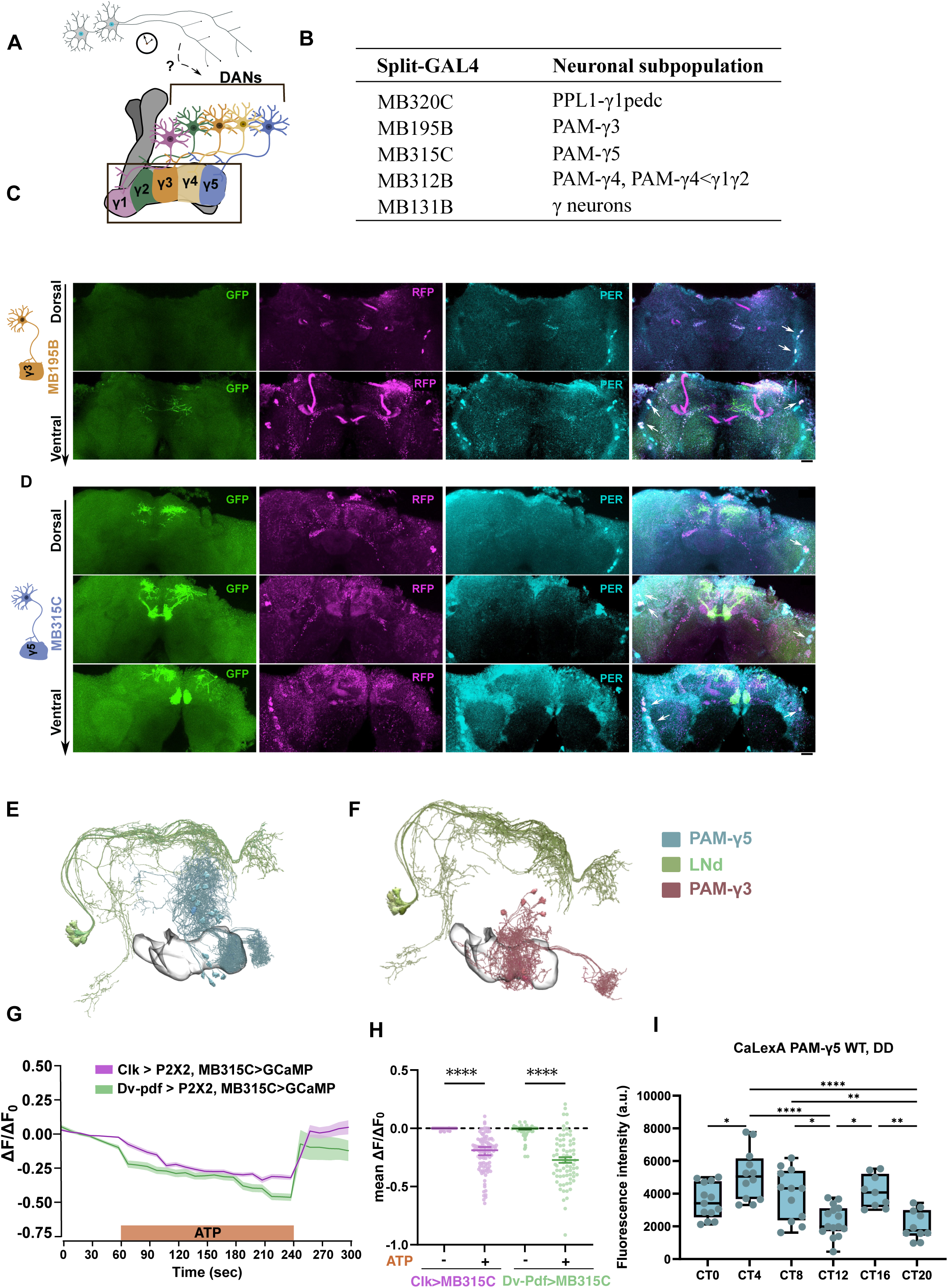
Functional and anatomical connectivity between the LNd clock neurons and the MB γ lobe via PAM-γ5 neurons. **A**. Schematic representation of the MB γ lobe compartments and their innervation by DA neurons. **B.** Split-GAL4 drivers used in retro-Tango experiments and neuronal subpopulations targeted. **C-D.** Representative images of retro-Tango-labeled neurons showing the postsynaptic DA neuron (green, expressing GFP), presynaptic partners (magenta, expressing mtdTomato), and clock neurons identified with anti-PER staining (cyan). White arrows indicate colocalization between the presynaptic partner and anti-PER staining, indicating the potential synapses with the LNds and l-LNvs (n= 20-30). Scale bar, 20um. Experiments were conducted at ZT20, coinciding with peak PER protein expression. **E and F**. NeuPrint skeleton representation of the PAM-γ5 neurons and the LNd (**E**) and l-LNvs (**F**), respectively. **G and H.** Analysis of functional connectivity between clock neurons and PAM-γ5 neurons using ATP-induced activation of P2X2 receptors. (**G**) The P2X2 channel was expressed using a pan-clock neuron driver (Clk-LexA) or DvPdf-LexA. GCaMP7s was expressed with a PAM-γ5-specific driver, MB315C. The average ΔF/F₀ of GCaMP7s fluorescence over time is plotted. The orange-shaded area indicates the ATP perfusion period, preceding and following periods represent baseline and HL3 washes, respectively. n=104 cells from 11 brains for the Clk>P2X2 group (pink trace). n=74 cells from 6 brains for the DvPdf>P2X2 group (green trace). The middle line is the mean, and the shadow represents the SEM**. H.** Quantification of mean ΔF/F0 during ATP perfusion compared to baseline levels. Mean ± SEM. Statistical comparisons were performed using a Kruskal–Wallis nonparametric test followed by Dunn’s multiple comparisons test. **I**. Calcium levels in PAM-γ5 neurons measured using the CaLexA reporter (n = 14–20) in DD at 6 timepoints. The center lines represent the median, box boundaries are the 25^th^ and 75th percentiles, and whiskers represent the minimum–maximum values. Statistical analysis was performed using two-way ANOVA with Dunnett’s correction for multiple comparisons. For clarity, only significant differences are indicated. Absence of a symbol denotes no significant difference. *p <0.05, **p<0.01, ***p<0.001, ****p<0.0001.

*Retro*-Tango tracing showed no evidence of a synaptic connection between circadian clock neurons and neurons labeled with the MB312B, MB320C, or MB131B driver. These results indicate that direct connectivity between clock neurons and PPL1, PAM-γ1, -γ2, -γ4, or γ-KCs is unlikely (Suppl. Figure 4A-C). In contrast, we found that presynaptic partners of PAM-γ3 and -γ5 neurons (labeled by MB195B and MB315C, respectively) included the LNd and l-LNv clock neurons (Figures 6C-D). In the fly brain connectome data (Schlegel et al., 2024), the LNds and PAM-γ5 neurons were found in close spatial proximity, displaying extensive contacts between their projections. In contrast, the l-LNvs did not appear in contact with PAM-γ5 neurons (Figure 6E, Supp. Figure 5A). Similarly, while PAM-γ3 neurons and the LNds were found in close proximity, the l-LNvs did not appear to be in contact with PAM-γ3 (Figure 6F, Supp. Figure 5B). Given that the projections of the l-LNvs terminate in distant, non-overlapping brain regions (Schubert et al., 2018), we conclude that the *retro*-Tango signals indicating the l-LNvs are presynaptic to PAM-γ5 and -γ3 neurons were likely artifacts.

Based on these results, we subsequently focused on PAM-γ5 neurons to test functional connectivity between clock neurons and DA neurons intermediary to the γ-KCs. Clock neurons were activated via the P2X2 ATP-gated cation channel, expressed under a pan-clock neuron driver, *Clk-LexA* (Cavanaugh et al., 2014). *UAS-GCaMP7s* driven by MB315C served as an indicator of intracellular calcium levels in PAM-γ5 neurons. Upon ATP perfusion, a significant decrease in GCaMP7s fluorescence intensity was observed (Figures 6G and H). This decrease was more pronounced when P2X2 was expressed with the *DvPdf-LexA* driver, which targets the LNds and s-LNvs (Figures 6G-H-). These findings support the anatomical data and demonstrate a functional inhibitory input from the LNds to PAM-γ5 neurons. This further suggests that input from clock neurons may rhythmically modulate the activity of PAM-γ5 neurons, thereby providing circadian control over the MB γ-lobe.

To test this, we monitored intracellular Ca2+ levels, as a proxy for neuronal activity, in PAM-γ5 neurons at 6 timepoints in DD using the CaLexA tool (Masuyama et al., 2012). The CaLexA signal exhibited a bimodal oscillatory pattern, with peaks in the middle of the subjective day (CT4) and the middle of the night (CT16) (Figure 6I). Importantly, prior research has shown that the LNds are most active from CT8 to 12 (Liang et al., 2016), which matches the valley in the calcium rhythms of PAM-γ5 neurons. Taken together, these data suggest that rhythmic inhibitory input from the LNds drives activity rhythms of PAM-γ5 neurons.

The temporal activity patterns of PAM-γ5 neurons closely resemble the cAMP and calcium oscillations (Machado Almeida et al., 2021) as well as the *Pka-C1* expression profile in γ-neurons (Figure 1F). This further suggests that excitatory input from PAM-γ5 neurons to MB γ-lobe drives γ-neurons’ activity rhythms, which in turn shape *Pka-C1* transcriptional rhythms. To test this hypothesis, we examined the effect of dopamine receptor silencing on *Pka-C1-luc* expression within γ-neurons. Four dopamine receptors (DopRs) – two D1-like receptors, a D2-like receptor, and a non-canonical receptor DopEcR –mediate DA signaling in flies. D1-like receptors, Dop1R1 (DUMB) and Dop1R2 (DAMB) are Gαs/Gαq-coupled and stimulate adenylyl cyclase (Feng et al., 1996; Gotzes et al., 1994; Sugamori et al., 1995). In contrast, D2-like receptor Dop2R is Gαi-coupled and inhibits adenylyl cyclase activity (Hearn et al., 2002). We therefore focused on the potential involvement of D1-like receptors Dop1R1 and Dop1R2. We knocked down each receptor in γ-neurons using two independent and validated RNAi lines (Hattori et al., 2017; Sun et al., 2018) (*Dop1R1*: BDSC line #31765 and #62193; *Dop1R2*: BDSC line #65997 and #26018). Knockdown of *Dop1R1* with either RNAi lines resulted in a significant reduction in *Pka-C1*-luc reporter signal and disrupted its rhythmicity (Figure 7A–B). In contrast, *Dop1R2* knockdown had no significant effect compared to controls.

**Figure 7.**
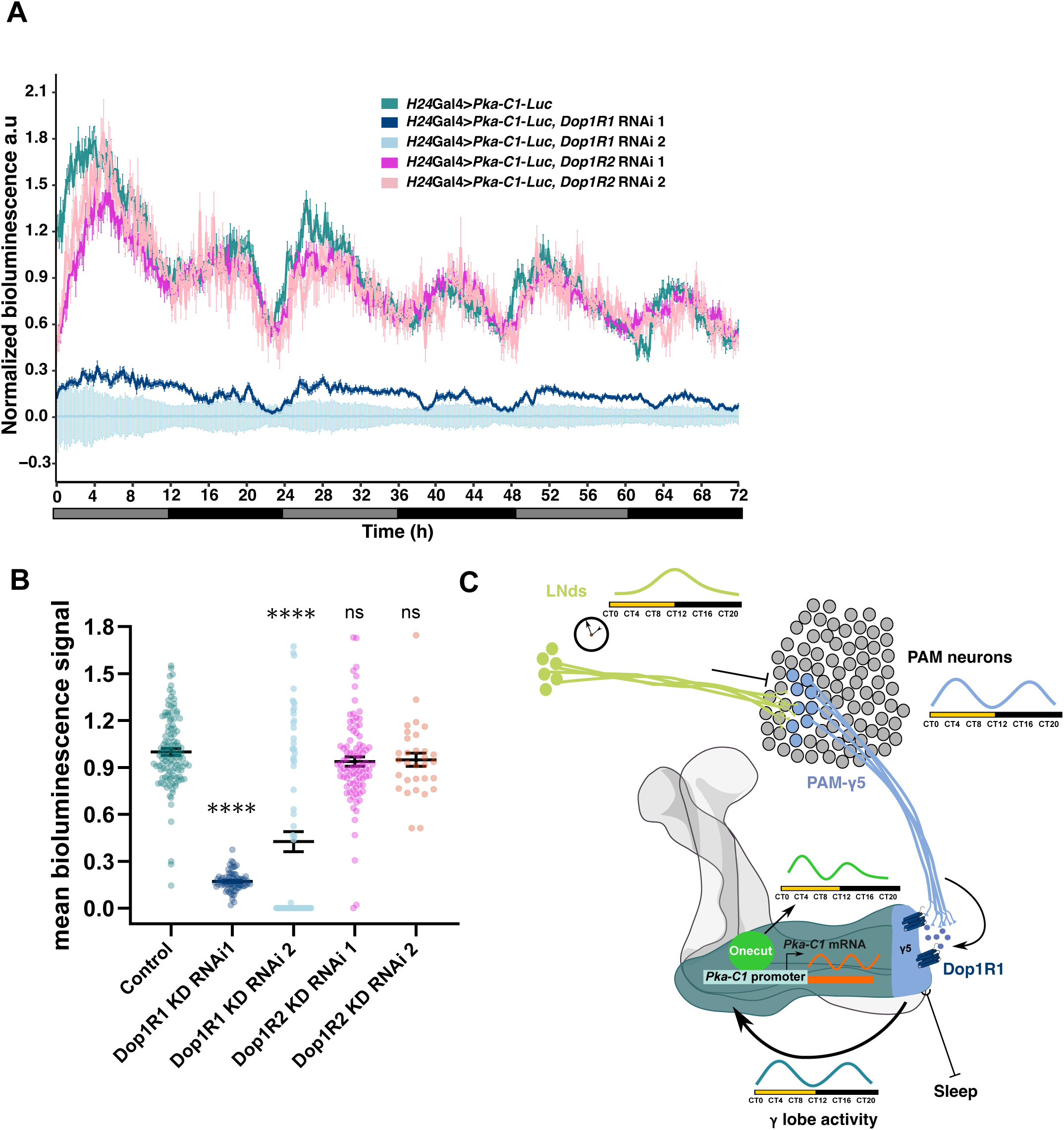
Clock neurons provide time-of-day signals to PAM-γ5 neurons, modulating their activity. **A and B.** Bioluminescence recordings of *Dop1R1* and *Dop1R2* knockdown (KD) flies in γ-neurons revealed a significant reduction in overall *Pka-C1*-luciferase signal in *Dop1R1* KD flies, but not in *Dop1R2* KD flies. This supports the involvement of dopamine signaling, likely originating from PAM-γ5 neurons, in modulating γ-neuron activity. In **A** bioluminescence recording for *control flies* (*H24 > Pka-C1-Luc,* aquamarine), *Dop1R1 KD* flies (dark blue: RNAi line #62193; light blue: RNAi line #31765), and Dop1R2 KD flies (pink: #26018; light pink: #65997). The middle line is the median, and the error bars represent ± the SEM. (**B**) Quantification of overall bioluminescence signal for each genotype. Data are shown as mean ± SEM. Statistical comparisons were performed using the Kruskal–Wallis nonparametric test followed by Dunn’s multiple comparisons test, comparing each KD genotype to the control (gray background). **C**. Proposed model: PAM-γ5 neurons serve as intermediaries between circadian clock neuron, specifically LNds, and MB γ-neurons, relaying time-of-day information via dopamine release. This signaling, mediated specifically through *Dop1R1*, regulates *Pka-C1* expression via Onecut and contributes to the circadian control of a wake-promoting circuit in the fly brain. *p <0.05, **p<0.01, ***p<0.001, ****p<0.0001. ns, not significant.

These findings provide strong evidence that PAM-γ5 neurons act as intermediaries between LNd clock neurons and MB γ-neurons, transmitting circadian signals via dopamine release. This dopamine release initiates a cascade within the γ-neurons: it drives cAMP oscillation via the excitatory DA receptor Dop1R1, which in turn regulates *Pka-C1* expression through Onecut transcription factor. The resulting PKA-C1 rhythms then generate intracellular calcium rhythms. Importantly, these activity rhythms reinforce PKA-C1 rhythms, creating a feedback loop between neuronal activity and transcriptional rhythms. This intricate feedback cycle promotes daytime wakefulness, likely through γ5 MB output neurons (MBONs) (Figure 7C). Our discovery that the Onecut–Pka-C1 pathway in the MB γ-neurons is a key component of this circuit significantly deepens our understanding of the sleep regulatory network.

## DISCUSSION

The MB is a crucial hub for information processing, integrating diverse sensory inputs and internal states to orchestrate complex behaviors. We demonstrate here that LNd clock neurons rhythmically inhibit PAM-γ5 DA neurons, which then rhythmically activate MB γ-KCs via excitatory Dop1R signaling. Resulting γ-neuron activity rhythms drive *Pka-C1* transcriptional rhythms through Onecut. Furthermore, these PKA-C1 rhythms reinforce neuronal activity rhythms, creating a feedback cycle between transcriptional and neural activity rhythms that promote daytime wakefulness. Our findings not only advance our understanding of sleep regulation but also broaden our insights into how temporal information is integrated into neural networks to shape daily patterns of behavior.

*In-silico* analysis and RNAi co-expressed with the *Pka-C1-luc* reporter led us to identify Onecut as a TF controlling rhythmic expression of *Pka-C1* in γ-neurons. This finding, however, does not exclude the involvement of other TFs in this process. The efficiency and specificity of gene silencing depend on the properties of the RNAi reagents, and available RNAis unfortunately did not yield high knockdown efficiency (Supp. Figure 2A). Nonetheless, the direct binding of Onecut to *Pka-C1* promoter, as shown by the ChIP experiments, and the increase in sleep upon *Onecut* knockdown in γ-neurons– the phenotype mirrors that of *Pka-C1* knockdown – provides supporting evidence for the role of Onecut in regulating *Pka-C1* expression.

Our strategy for mutating Onecut binding regions in the *Pka-C1* promoter resulted in various indels near the predicted binding site in either the endogenous gene or the transgene promoter (Figure 5 and Supp. Figure 3). This approach has certain limitations, including the inability to generate identical mutations in both contexts. Nevertheless, the finding that mutations in either the endogenous gene or the transgene similarly disrupt *Pka-C1*-*luc* rhythms revealed a crucial feedback loop from neuronal activity rhythms to *Pka-C1* transcriptional rhythms. Onecut is a transcription factor essential for neuronal development and function in *Drosophila*, with roles in differentiation, axon guidance, and brain activity. It binds DNA via two domains, the CUT domain and the homeodomain, which cooperate to regulate transcription. Importantly, a prior study has shown that mutations in the 5′ flanking region of the core ATTG binding motif reduce Onecut binding efficiency to a greater extent than mutations within the ATTG motif (Nguyen et al., 2000). Our ChIP-qPCR experiments, which revealed significant Onecut binding in the regions flanking the ATTG motif, further support this prior finding. Together, these results indicate that our CRISPR mutations likely reduce Onecut binding efficiency.

*Onecut* mRNA and protein accumulate rhythmically in the MB KCs (Figure 4A-C) (Machado Almeida et al., 2021). Onecut protein oscillations are in phase with *Pka-C1* transcription (Figure 4A-C). Given the feedback regulation between PKA-C1 rhythms and neural activity rhythms, it is plausible that Onecut expression rhythms are regulated by neuronal activity. Consistent with this, there is evidence that Onecut’s transcriptional activity is triggered by translational cascades activated by sensory inputs such as olfaction (Jafari et al., 2012), vision (Vassalli et al., 2021), and hearing (Keder et al., 2020). In mammals, phosphorylation of Onecut’s homolog, HNF-6, by PKA is required for G6Pase gene expression, and PKA phosphorylation enhances HNF-6 binding to its target promoter (Streeper et al., 2001). This suggests that PKA-mediated phosphorylation may also be involved in the regulation of Onecut in *Drosophila*. Additionally, Onecut binding sites are highly enriched among genes rhythmically expressed in the MB (Machado Almeida et al., 2021). It is also among the transcription factors with the highest target enrichment in the adult Drosophila CNS (Davie et al., 2018). Therefore, Onecut likely regulates not only Pka-C1 transcription but also other genes that control MB function.

We establish here an indirect pathway, the LNd – PAM-γ5 – MB γ lobe circuit, through which circadian pacemaker neurons promote daytime wakefulness (Figure 7C). Our findings are consistent with previous studies demonstrating the wake-promoting roles of PAM-γ5 and MB γ5 subdomain, mediated by DA signaling via Dop1R1 in the γ-KCs (Artiushin & Sehgal, 2017; Driscoll et al., 2021). Inhibition of PAM-γ5 neurons by the LNds serves as a circadian mechanism to suppress arousal signals during the rest phase, linking the molecular clock to sleep-wake regulation. The LNds are a heterogeneous group composed of at least three molecularly distinct subtypes, including neurons that express ion transport peptide (ITP), short neuropeptide F (sNPF), and neuropeptide F (NPF). Both sNPF and NPF signal through Gi/o-coupled receptors (sNPFR1 and NPFR1, respectively), which are known to reduce intracellular cAMP levels and suppress neuronal excitability (Garczynski et al., 2002; Liu et al., n.d.; Shang et al., 2013). Additionally, both neuropeptides are involved in sleep-wake modulation(Chung et al., 2017; Shang et al., 2013). Therefore, although the precise mechanism by which the LNds inhibit PAM-γ5 DA neurons remains unknown, signaling via sNPF and/or NPF may be a plausible mechanism.

PKA signaling in the MB is essential for memory and associative learning, suggesting that this newly discovered circuit may be a shared mechanism linking circadian modulation of sleep and memory—two processes thought to be interconnected. In particular, the γ lobe is required for memory acquisition and formation of short-term memory immediately after associative learning (Qin et al., 2012; Swinderen, 2009; Zars et al., 2000). Importantly, Dop1R1 is associated with memory formation, whereas Dop1R2 is implicated in active forgetting (Handler et al., 2019). Thus, the dependence of *Pka-C1* transcriptional rhythms and sleep modulation on Dop1R1 provides insight into a shared pathway that may act as a molecular filter between sleep and memory formation.

Collectively, this work uncovers a pathway connecting circadian inputs to MB, integrating molecular and neuronal mechanisms to regulate complex behaviors. It highlights the intricate interplay between the circadian system and brain regions governing sleep and arousal. Our findings not only advance our knowledge of the pathways underlying rhythmic gene expression and neuronal activity but also underscore the broader significance of circadian regulation in shaping complex, state-dependent behaviors.

## MATERIALS and METHODS

All the reagents and antibodies are described in Supplementary Table 1. Sequences of primers and gRNAs are listed in Supplementary Table 2.

### Fly strains

All *Drosophila melanogaster* strains were maintained on standard cornmeal-agar food at 18 °C or 25 °C in a 12-h light/ 12-h dark cycle and controlled humidity. The lines *w^1118^* and *per^0^* were previously described (Grima et al., 2004). Detailed information on the strains used in the present study is provided in Supplementary Table 3.

### Immunohistochemistry and microscopy

The CaLexA (calcium-dependent nuclear import of LexA) technique was employed to monitor neuronal activity in the PAM-γ5 dopaminergic subpopulation (Masuyama et al., 2012). GFP expression in PAM-γ5 neurons was visualized via whole-mount brain immunofluorescence with anti-GFP and anti-TH antibodies. To assess Onecut protein levels, the cell bodies of γ-KCs labeled with GFP using the γ-neuron-specific H24-Gal4 driver were anaylzed. Quantification was performed in regions where GFP and anti-Onecut immunoreactivity colocalized. In both experiments, flies aged 5–6 days were decapitated at six timepoints (ZT/CT 0, 4, 8, 12, 16, and 20) following 3 days of LD entrainment. For experiments in DD, brains were collected on DD2 after the LD entrainment. Neural circuit tracing was conducted using the *retro*-Tango tool, which utilizes ligand-receptor interactions at synapses to trigger a signaling cascade resulting in the expression of mtdTomato in presynaptic neurons and GFP in postsynaptic neurons (Sorkaç et al., 2023). GFP expression in PAM-subpopulations was driven by specific split-GAL4s. Visualization was performed using whole-mount brain immunofluorescence with anti-GFP, anti-RFP, and anti-PER. 5–7-day-old flies were decapitated at ZT20, coinciding with peak TIM and PER expression.

All the imaging experiments were conducted at least in duplicate and for a sample size of at least 15-20 flies per replicate. The preparation procedure was consistent across experiments, except for the primary and secondary antibodies used (details and concentrations are listed in Supplementary Table 1). Flies were decapitated and heads were fixed in 4% paraformaldehyde containing 0.3% Triton X-100 for 1 hour on ice. Samples were washed twice briefly and twice for 20 minutes with PBST-0.3 (PBS, 0.3% Triton X-100). The head cuticles were partially removed before blocking in a solution containing 5% normal goat serum in PBST-0.3 for 1 hour at room temperature. Samples were then incubated with primary antibodies for 48 hours at 4°C, followed by washing (2 quick washes and 2 washes of 20 minutes each) and incubation with secondary antibodies for 24 hours at 4°C. After the secondary antibody staining, heads were washed. Then, the remaining cuticle was removed, and brains were mounted on slides using Vectashield mounting medium (Vector Laboratories). Imaging was performed using a Nikon AX confocal microscope.

### Luciferase reporter constructs

To generate transgenic flies expressing a luciferase reporter under the control of the *PkaC1* promoter with or without the *PkaC1* 3’UTR, a plasmid designed by Dr. Rafael Koch and Dr. Pedro Machado Almeida was utilized as the backbone. The backbone plasmid was pre-assembled with several elements, excluding the specific promoter and 3’UTR sequences of interest. These elements included a codon-optimized coding sequence (CDS) for firefly luciferase, an FRT-STOP-FRT cassette, the p’10 3’UTR sequence, gypsy insulator sequences, and an attB phiC31 integrase site for targeted genomic integration. To obtain *Pka-C1-luc* and *Pka-C1-luc-3’UTR* plasmids, the original promoter in the backbone was replaced with the *PkaC1* promoter sequence. For the second construct, the p10 3’UTR sequence was further replaced with the *PkaC1* 3’UTR sequence.

#### Backbone origin and details

The backbone was derived from the pBID-13xLexA-Gateway-p10 3’UTR (a gift from Dr. Brian McCabe), which contains gypsy insulator sequences. The FRT-STOP-FRT cassette was PCR-amplified from the pUAS-mCD8-GFP plasmid (Addgene #24385), with terminal modifications to remove restriction sites. The codon-optimized firefly luciferase (Fluc) CDS for *Drosophila* was obtained from the pCa4B2G.UAS-FLuc plasmid (Markstein et al., 2008). ATG codon was removed from the Fluc sequence to minimize miss-expression. It was placed between the promoter and the first FRT sequence, allowing the Fluc coding sequence to be in frame with the ATG codon only after the FRT recombination. These components were synthesized and assembled into the backbone by Custom DNA Constructs Company to create a functional plasmid.

#### Pka-C1-luc plasmid construction

From the backbone plasmic, the *Act5* promoter was removed using AgeI and EcoRI restriction enzymes. The linearized vector was purified by gel extraction. Using a Gibson assembly kit (NEB; see Table 3 for references) a PCR-amplified *Pka-C1* promoter sequence was assembled into the digested backbone vector using the Gibson assembly method. The *Pka-C1* promoter with 20 bp ends overlapping the insert site of the vector was amplified by PCR using a Q5 polymerase from genomic DNA of *w^1118^*flies, covering the region from −2107 bp up to the start codon. Gibson assembly was performed using the Gibson Assembly Clonig Kit (NEB). The amplified promoter sequence and successful assembly was confirmed by Sanger sequencing.

#### Pka-C1-luc-3’UTR plasmid construction

The p’10 3UTR sequence in the backbone plasmid was removed using the Acc65I and KflI enzymes. The *Pka-C1* 3’UTR sequence was amplified by PCR from the BAC CHORI-322-172P9, using primers designed to incorporate restriction sites for Acc65I and KflI enzymes. The amplified *Pka-C1* 3’UTR sequence was purified and confirmed by Sanger sequencing before being digested with Acc65I and KflI. The linearized vector was purified by gel extraction, and the digested PkaC1 3’UTR sequence was then ligated into the vector using T4 DNA ligase. The complete sequence of the assembled vector was confirmed by Sanger sequencing.

#### Amplification of the constructs

To amplify both constructs, *E. coli* competent cells were transformed and plated. To minimize the frequency of recombination events, 20% sucrose was added to the LB medium. Positive colonies were picked, cultured, and the amplified plasmids were verified by Sanger sequencing. The constructs were then sent to BestGene Inc. (California, USA) for embryo injections and the creation of transgenic lines. Both vectors were integrated into the attP40 site.

### Mutagenesis using the CRISPR/Cas9 system

To generate mutants in the *Pka-C1* promoter, we first generated two transgenic fly lines, each carrying a different UAS-gRNA targeting the Onecut binding site within the *Pka-C1* promoter. The two gRNA sequences were designed to be closely spaced and located near the Onecut binding site to maximize targeting efficiency. The gRNA sequences (Supplementary Table 2) were designed using the CRISPR Target Finder web tool (http://targetfinder.flycrispr.neuro.brown.edu/), which evaluates target specificity to minimize off-target effects. The gRNAs were synthesized by VectorBuilder and inserted individually into the pCFD6 vector backbone (Addgene #73915). The sequence of the constructs was confirmed by Sanger sequencing. The finalized constructs were sent to BestGene Inc. (California, USA) for embryo injections and the generation of transgenic lines. Both constructs were integrated into the attP154 landing site on the 3rd chromosome of the *Drosophila melanogaster* genome. To generate mutations, males of the UAS-gRNA transgenic lines were crossed with females carrying the *nos-GAL4* driver, which is expressed in the female germline. Screening for indels in the target region was performed using qPCR with Custom TaqMan® Gene Expression Assay probes designed to specifically detect and discriminate between both the mutated and wild-type alleles. Seven distinct mutant lines were identified and further validated by Sanger sequencing. Two lines were selected for the following studies.

### Live bioluminescence measurements

For each experiment, each well of the white 96-well half-area plates (CORNING #3693) were filled with 100 µL of standard behavioral fly food (5% sucrose and 2% Bacto agar), overlaid with 25 µL of standard behavioral food containing 15 mM D-luciferin dissolved in water. Prior to bioluminescence recording, flies were entrained for three complete cycles of 12h/12 h LD. Flies were kept at 25°C during the whole process, with the exception of the TrpA1 experiments. Flies expressing TrpA1 were raised at 18°C, and the bioluminescence recording was performed at 29°C to activate the TrpA1 channel. To minimize signal fluctuations due to habituation, flies were placed in the prepared plate one day before the recording. Each fly was individually positioned in a well, following a zig-zag pattern to reduce signal crosstalk and include wells for background signal measurements, which were subtracted during analysis. The plates were sealed with Ultra Clear sealing films (Axygen, cat. no. UC-500) punctured twice with a needle to allow airflow. Bioluminescence recording was performed using a plate reader (LUMIstar Omega, BMG Labtech) for 74 hours under DD, starting 2 hours before CT0 (i.e., ZT20 in the last LD cycle). The first 2 hours of data were excluded from analysis to exclude the data during acclimatization. The recording settings included an emission gain of 3600, an integration time of 4.7 seconds per read, and a total of 555 reads per experiment. For recordings of PAM-γ5 neurons, where the cell population was smaller, the emission gain was increased to 4096. With these settings, bioluminescence signals from up to 48 flies were recorded simultaneously in each experiment. Each experiment was performed in at least duplicate, with a sample size of 20–48 flies per replicate, including both males and females.

### Bioluminescence data analysis

All analyses were performed using RStudio (v.2024.09.1). Since the data were collected with an integration time of 4.7 seconds, the initial transformation involved converting the values to counts per second and subtracting the background signal calculated from empty wells. Each fly’s bioluminescence counts were normalized to its total mean. The analysis methodology was based on a modified version of Martin-Burgos et al. (2022) (Martin-Burgos et al., 2022). Briefly, a discrete wavelet transform (DWT) was applied to each time series to detrend the data and determine rhythm phase, using the wavelets (v. 0.3-0.2) R package as described by Leise and Harrington (2011) and Leise (2017). The la12 filter was applied to 30-minute median-binned data, where medians were used to reduce the impact of large outliers. After detrending, the statistical significance of circadian rhythms for individual flies was assessed using maximum entropy spectral analysis (MESA) power value. These analyses were conducted over 3-day windows, using 120-minute median binning to minimize correlated high-frequency noise and outliers. The significance threshold was set to a power value >0.2. This algorithm was also used to calculate the period of the bioluminescence rhythm for each individual fly. Graphs for bioluminescence signals were generated in R using the ggplot2 (v3.1.0) package, with data represented as median ± standard error. Statistical comparisons and plotting of the rhythmicity index (RI) and the percentage of *Pka-C1-Luc* rhythmic flies were conducted using GraphPad Prism (v.10.4.0). Depending on whether the data followed a normal distribution, either a t-test or a Mann-Whitney test was used for statistical analysis in the RI, and a Chi-square test was used to determine the significance of *Pka-C1-Luc* rhythmic flies.

For Dop1R1 bioluminescence data analysis, to account for the decrease in signal intensity, we adapted the analysis by focusing on comparing overall signal levels to the control group. Raw luminescence values were first converted to counts per second, and background signal—calculated from empty wells—was subtracted. A normalization factor was then computed based on the overall mean bioluminescence signal across all control flies throughout the day. This factor was used to normalize the individual bioluminescence traces of both *Dop1R1* knockdown and control flies. Bioluminescence traces were visualized in R using the ggplot2 package (v3.1.0), with data represented as median ± standard error. To statistically compare the overall mean signal per genotype to the control, we used GraphPad Prism (v10.4.0) and conducted a Kruskal–Wallis test followed by Dunn’s multiple comparisons test, using the control group as the reference.

### P2X2 Activation and GCaMP imaging

Adult male and female flies of 5-7 days old were entrained for at least 3 days in a (12h:12h LD) cycle before functional imaging experiments. The experiments were conducted following the protocol is described in Barber AF. et al. (2021) (Barber et al., 2021). Briefly, flies were anesthetized on ice and dissected in hemolymph-like saline (HL3)(Yao et al., 2012). Imaging experiments were performed in a perfusion chamber (Ibidi, µ-Slide I Luer 3D, #87176). The slide was first coated with a thin layer of collagen and polymerized at 37°C for 2 hours to stabilize the brain before the experiment. The brain was then placed in the chamber in a small bath of HL3. Solutions were perfused over the brain at a rate of ∼5 mL/min with a gravity-fed 8-Channel Pinch Valves automated perfusion system (Bioscience Tools). After 1 min of baseline GCaMP7s imaging, ATP was delivered to the chamber by switching perfusion flow from the channel containing HL3 to another channel containing 2.5 mM ATP (Sigma-Aldrich, #A2383-5G) in HL3, pH 7.1. ATP was perfused for 2.5 min, followed by a 1-min washout with HL3. GCaMP7s calcium imaging was performed on a Nikon AX confocal microscope. Twelve-bit images were acquired with a 4x air objective with 10X zoom at 256 x 256-pixel resolution. Z-stacks were acquired every 10 s.

Image processing and measurement of fluorescence intensity were performed in FIJI as previously described (Barber et al., 2021). A sum-intensity Z-projection of each time step was used for analysis, and the StackReg FIJI plugin was used to correct for small x-y movements over time in the sum-projected image. Regions of interest (ROIs) were manually drawn to encompass individual GCaMP-positive cell bodies, and mean fluorescence intensities were measured from each ROI at each time point. For each cell, fluorescence traces over time were normalized using this equation: ΔF/F = (Fn-F0)/F0, where Fn is the fluorescence intensity recorded at time point n, and F0 is the average fluorescence value during the 30-s baseline preceding ATP application. The mean ΔF/F was calculated based on the average ΔF/F during the ATP stimulation period. Brains containing cells with unstable baselines were excluded from the analysis. Statistical comparisons between two groups were performed using Student’s *t*-test.

### ChIP-qPCR

ChIP was performed as described previously (Zhou et al., 2015) with minor modifications, on male and female *w^1118^* flies. Briefly, approximately 2000 7-day-old flies per sample were collected 4 h after lights-on (ZT4), the peak time of Onecut protein expression in LD, and stored at −80 °C until processed. Heads were collected using metal sieves. Approximately 1 ml of frozen fly heads (∼20 ml of flies) was ground on dry ice using a pre-chilled 7-ml homogenizer (80 strokes). The homogenized material was immediately resuspended in 5 ml freshly prepared XIP homogenization buffer containing 50 mM HEPES-KOH pH 8.0, 140 mM NaCl, 1 mM EDTA pH 8.5, 0.5 mM EGTA pH 8.0, 0.4% Igepal CA-630, 0.2% Triton X-100, 1% formaldehyde, 1 mM PMSF, 1 mM Na₃VO₄ and Protease inhibitor cocktail (Roche 11 873 580 001)). Immediately after resuspension, samples were gently homogenized for 10 min at room temperature with occasional vortexing. Cross-linking was quenched by adding glycine (final concentration 0.125 M) and continuing the homogenization for 5 min at RT. The homogenate was filtered through 100 μm nylon mesh and centrifuged at 1500 × g for 5 min. The pellet was washed and resuspended in wash buffer (20 mM Tris-HCl pH 7.5, 150 mM NaCl, 1 mM EDTA pH 8.5, 0.5 mM EGTA pH 8.0, 0.1% Igepal CA-630, 0.1% Triton X-100, 0.5 mM PMSF, Protease inhibitor cocktail (Roche 11 873 580 001) and 1 mM Na₃VO₄). Cross-linked nuclei were resuspended in sonication buffer (10% glycerol, 1% Triton X-100, 0.4% sodium deoxycholate, 0.1% SDS, 10 mM Tris-HCl pH 7.5, 150 mM NaCl, 1 mM EDTA, 0.5 mM EGTA, 0.5 mM PMSF, Protease inhibitor cocktail (Roche 11 873 580 001), 1 mM Na₃VO₄). Samples were sonicated using Branson Ultrasonic digital sonifier 450 (30 × 5 s at 4–5 W with 25 s rest on ice bath). After centrifugation at 25,000 × g for 10 min at 4 °C, the supernatant was collected, and protein concentration was determined by Pierce™ BCA Protein Assay Kits (#23225). For immunoprecipitation, 500 μg of protein–DNA complexes were incubated overnight at 4 °C with 3 μl of anti-Onecut antibody (Supplementary Table 1). Dynabeads Protein A/G (30–50 μl) were blocked with blocking buffer (0.1 μg/μl sonicated salmon sperm DNA, 5 μg/μl BSA in immunoprecipitation buffer). The IP buffer contained 1% Triton X-100, 0.01% SDS, 10 mM Tris-HCl pH 7.5, 150 mM NaCl, 1 mM EDTA, 0.5 mM EGTA, 0.5 mM PMSF, Protease inhibitor cocktail (Roche 11 873 580 001), and 1 mM Na₃VO₄. Beads were incubated with chromatin– antibody complexes for 2 h at 4 °C. After binding, immunocomplexes were sequentially washed with: (1) IP buffer (twice), (2) Low-salt buffer (same as IP buffer but with 150 mM NaCl), (3) High-salt buffer (same as IP buffer but with 500 mM NaCl), (4) LiCl buffer (10 mM Tris-HCl pH 7.5, 1% Igepal CA-630, 1% DOC, 250 mM LiCl, 1 mM EDTA), and (5) TE buffer (twice). Elution was performed with elution buffer (1% SDS, 100 mM NaHCO₃ in water) at 65 °C for 15 min, twice. Eluates were pooled (100 μl total), treated with RNase A (50 μg/ml, 30 min, 37 °C), followed by proteinase K (1 μg/μl, 6 h to overnight, 37 °C), and crosslinks were reversed overnight at 65 °C. DNA was extracted by phenol–chloroform and precipitated in ethanol with glycogen (0.5 μl of 20 μg/μl) and salmon sperm DNA (0.5 μg). After ethanol washes and drying, DNA was resuspended in 50 μl TE buffer. The qPCR was performed using the DNA samples obtained, which were amplified using the Thermo Fisher QuantStudio 5 system. Enrichment was quantified using a standard curve generated from serial dilutions of genomic DNA, and fold change was calculated. Primer sequences and regions used are listed in Supplementary Table 2.

### Sleep assay and analysis

The sleep assay was conducted using the Drosophila Activity Monitoring (DAM) system (Trikinetics) as described previously (Shaw et al., 2000). Briefly, 2-to 5-day-old male or virgin female flies were individually placed in 65 × 5 mm glass tubes containing 5% agarose and 2% sucrose as a food source. Flies were entrained under 12h/12h LD cycles for three days, and sleep data were collected during the subsequent three days of LD. Locomotor activity was recorded in 1-minute bins, and sleep was defined as periods of inactivity lasting 5 minutes or longer. All sleep assays were conducted at 25°C. Data analysis was performed in MATLAB (MathWorks, version R2022b, build 9.13.0.2049777) using SCAMP (v3.0, Vecsey Laboratory) following the instruction manual. Statistical comparisons were performed on the raw data obtained using SCAMP. Two-way ANOVA followed by Šìdák’s correction or Mann–Whitney tests with Šìdák’s correction were applied for multiple comparisons, depending on whether the data followed a normal distribution. Statistical analysis and plots were generated in GraphPad Prism (v10.4.0).

### Image analysis

Confocal Z-stacks were analyzed using ImageJ/Fiji software (v2.1.0/1.53c) (Schindelin et al., 2009). CaLexA GFP fluorescence intensities in individual PAM-γ5 neurons were quantified from sum-slice Z-projections, generated every five stacks, within the area defined by the TH signal. Then, the mean signal per brain was calculated. For *retro*-Tango experiments, colocalization between RFP was assessed. Anti-Onecut and GFP fluorescence intensities in γ neurons were quantified from sum-slice Z-projections generated every five stacks, within the area defined by the GFP signal. The background signal was subtracted for both, and the anti-Onecut signal was normalized with the GFP signal. Then, the average anti-Onecut and GFP signal was calculated.

### Statistics and reproducibility

GraphPad Prism (v10.4.0) was used for statistical analysis and data plotting unless otherwise indicated. Normally distributed data were compared using parametric tests and non-normally distributed data were analyzed using nonparametric tests. Sleep data were analyzed using Two-way ANOVA followed by Šìdák’s correction or Mann–Whitney tests with Šìdák’s correction.

Statistical comparison of fluorescence intensities was performed using the ordinary one-way ANOVA with Bonferroni’s multiple comparison test when data were normally distributed, and the Kruskal–Wallis with Dunn’s multiple comparison test was used to compare non-normally distributed data. Significant values in all figures are: *p < 0.05, **p < 0.01. ***p < 0.001, and ****p < 0.0001. ns indicates not significant. All the experiments were repeated at least twice.

## ACKNOWLEDGMENTS

We thank Donggen Luo and the Bloomington *Drosophila* Stock Center for providing fly stocks and Brian McCabe for the plasmids. We are grateful to our lab members for their daily help and valuable discussions. This research was supported by grants from the Swiss National Science Foundation (189169 and 1000329), Georges and Antoine Claraz Foundation, Fondation Schmidheiny, and Société Académique de Genève.

## AUTHOR CONTRIBUTIONS

BL and EN conceptualized the study. BL designed experiments, performed experiments, and analyzed data. BL produced visualizations of the results and made figures. BL and EN wrote the manuscript.

## SUPPLEMENTARY INFORMATION

### Supplementary Figures

**Supplementary Figure 1.**
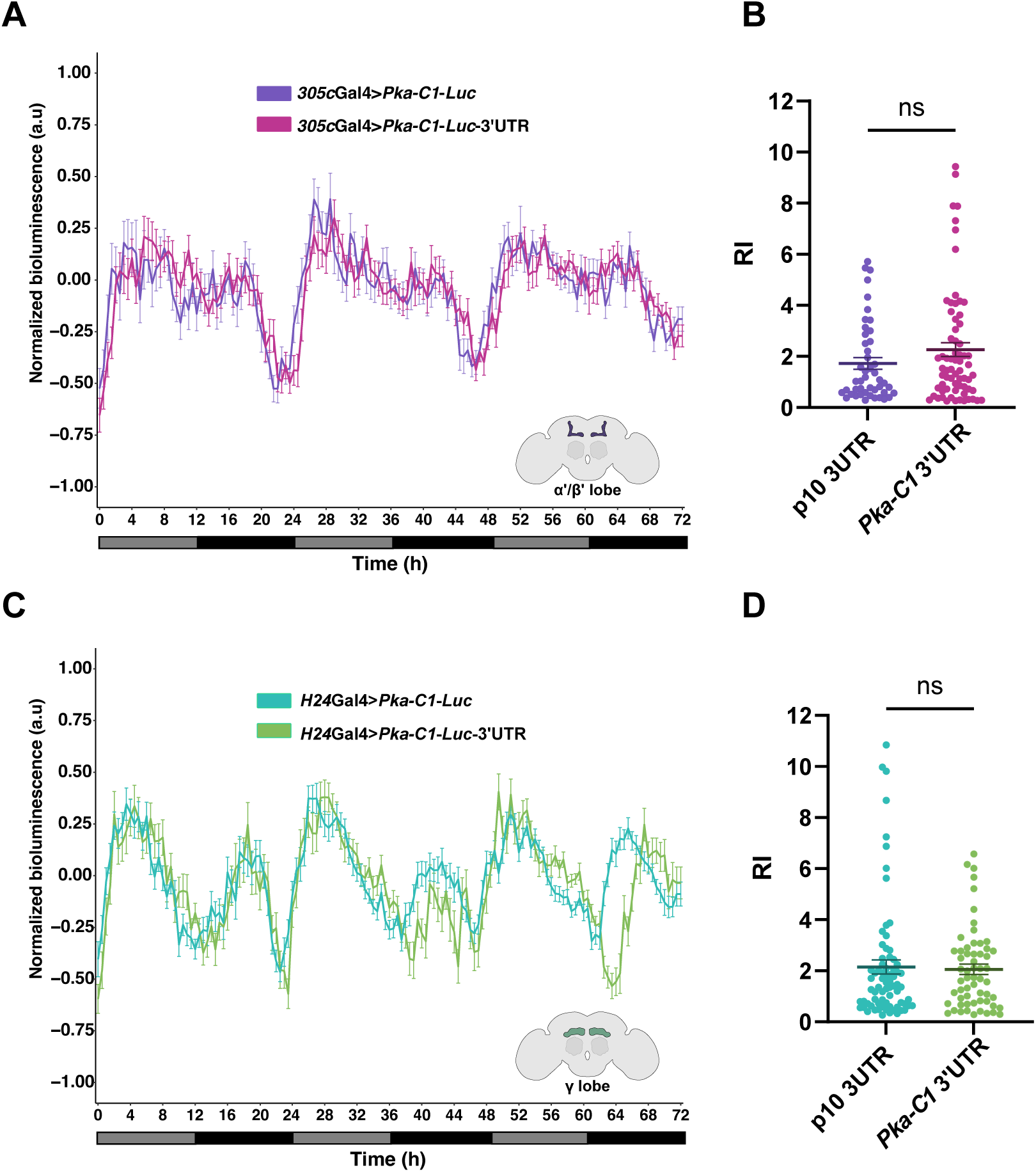
*Pka-C1* 3’UTR does not increase the strength of *Pka-C1* rhythms in α′/β- and γ-KCs. **A and C.** Bioluminescence recordings of *Pka-C1-luc* and *Pka-C1-luc-*3’UTR reporter expressed in α′/β′-KCs (n=98) (**A**) or γ-KCs (n=87) (**C**). The middle line is the median, and the error bars represent the SEM. **B and D.** RI of reporter bioluminescence rhythms. The center lines and the error bars represent the mean ± SEM. Statistical comparisons were performed using a t-test or Mann-Whitney test, depending on the data distribution. *p <0.05, **p<0.01, ***p<0.001, ****p<0.0001. ns, not significant.

**Supplementary Figure 2.**
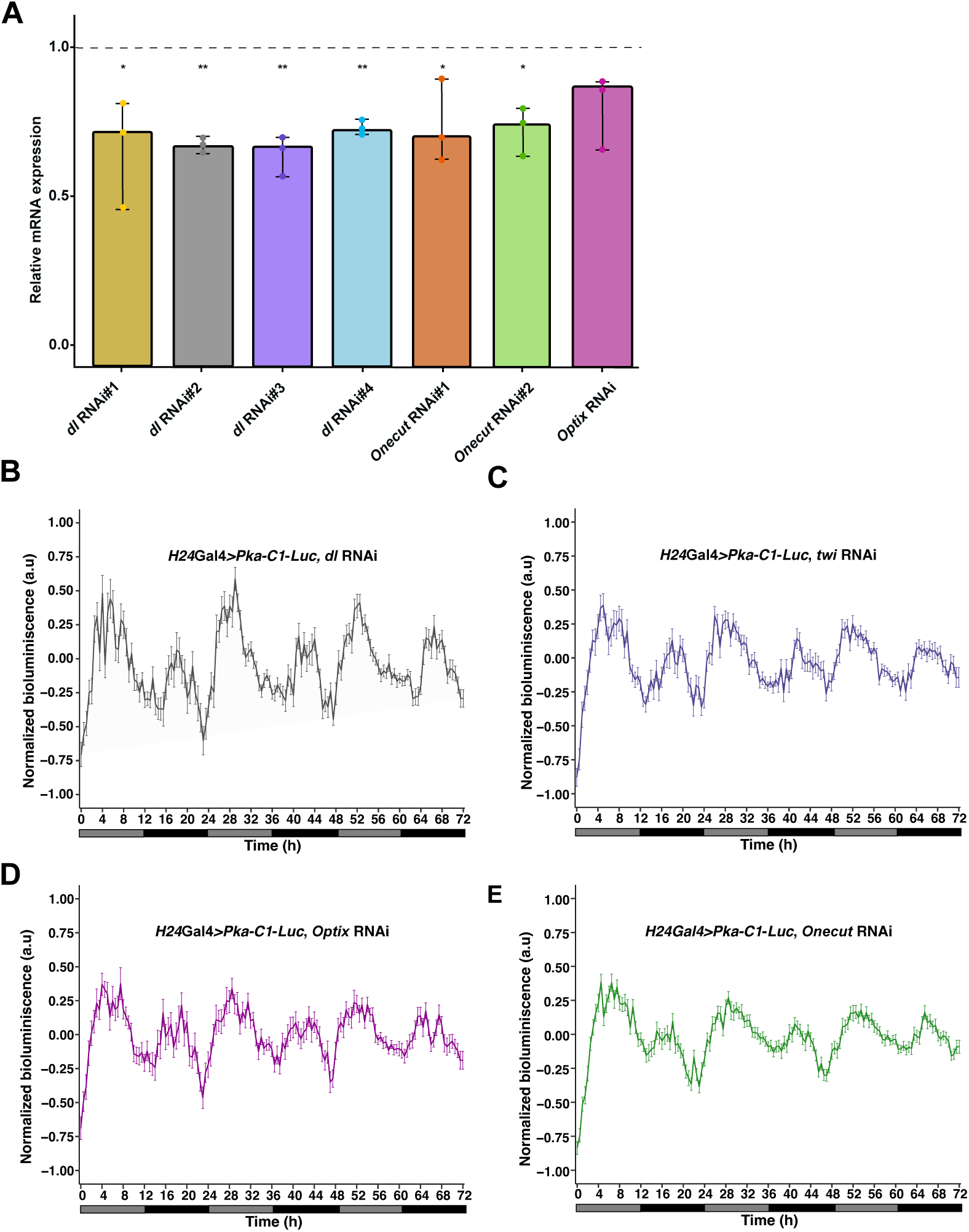
Knockdown efficiency of TF RNAi lines and bioluminescence recording of KD lines. **A.** The different RNAi lines were driven with a pan-neuronal Gal4 driver (*elav*-Gal4), and expression levels of the target mRNAs were quantified using qPCR. The value indicates the relative mRNA levels of the target gene compared to those in the control (*elav*-Gal4/+) flies. Bar plots show the mean, minimum, and maximum. Each dot represents an individual replicate (24 flies per group). The Kruskal–Wallis nonparametric test followed by Dunn’s multiple comparisons test was used to compare values to the control. **B-E.** *H24>Pka-C1-luc* bioluminescence recordings of flies with KD of each of the four TFs. RNAi against *dorsal* (n= 63) (**B**), *twist* (n= 90) (**C**) *Optix* (n= 68) (**D**), and *onecut* (n= 105) (**E**). The middle line is the median, and the error bars represent ± the SEM. *p <0.05, **p<0.01, ***p<0.001, ****p<0.0001. ns, not significant.

**Supplementary Figure 3.**
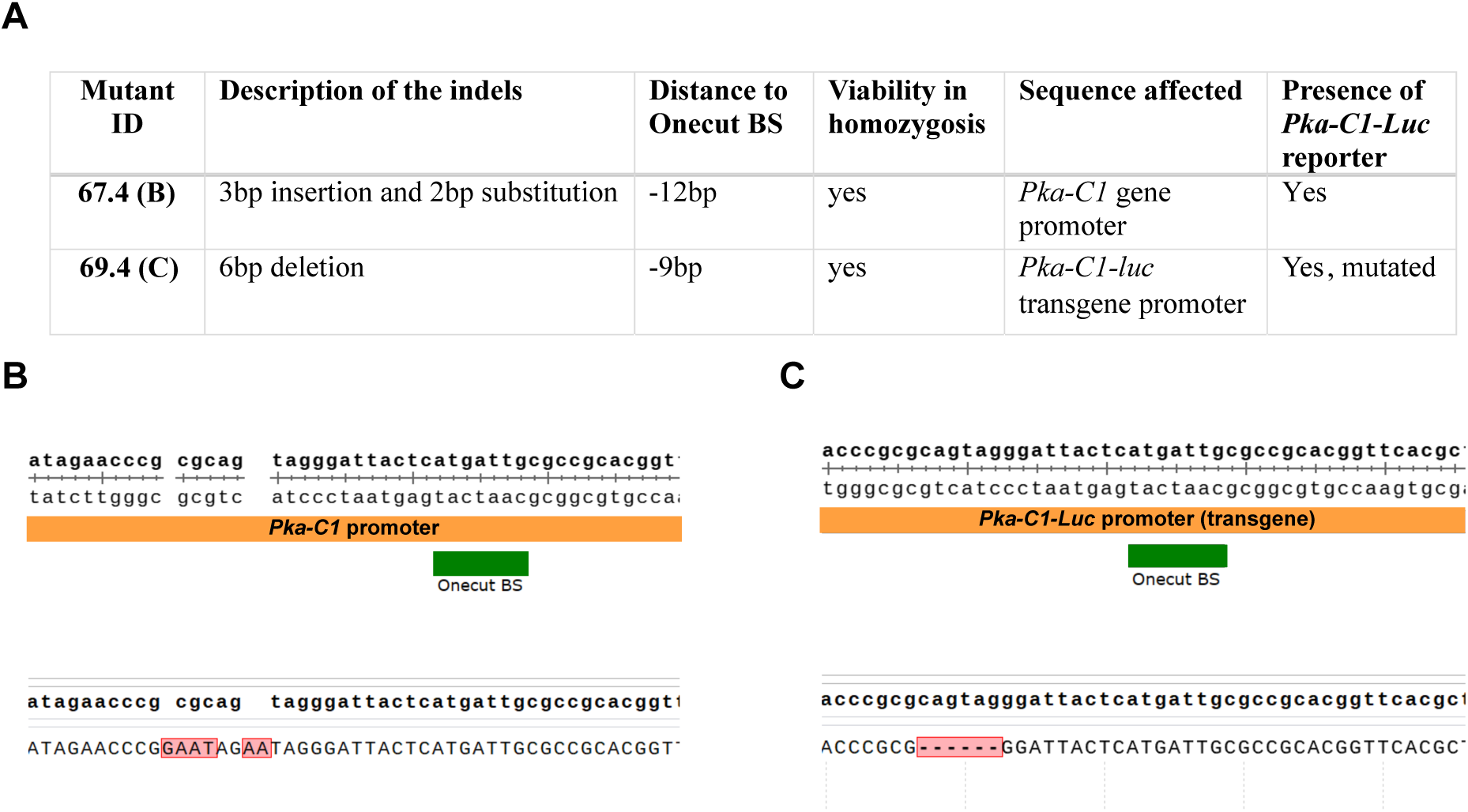
CRISPR mutant lines. **A.** Characteristics of the two CRISPR mutant lines obtained. BS, binding site. **B and C.** Schematic representation of the mutations in the *Pka-C1* gene in the line 67.4 (**B**) and *Pka-C1-luc* promoter in the line 69.4 (**C**).

**Supplementary Figure 4.**
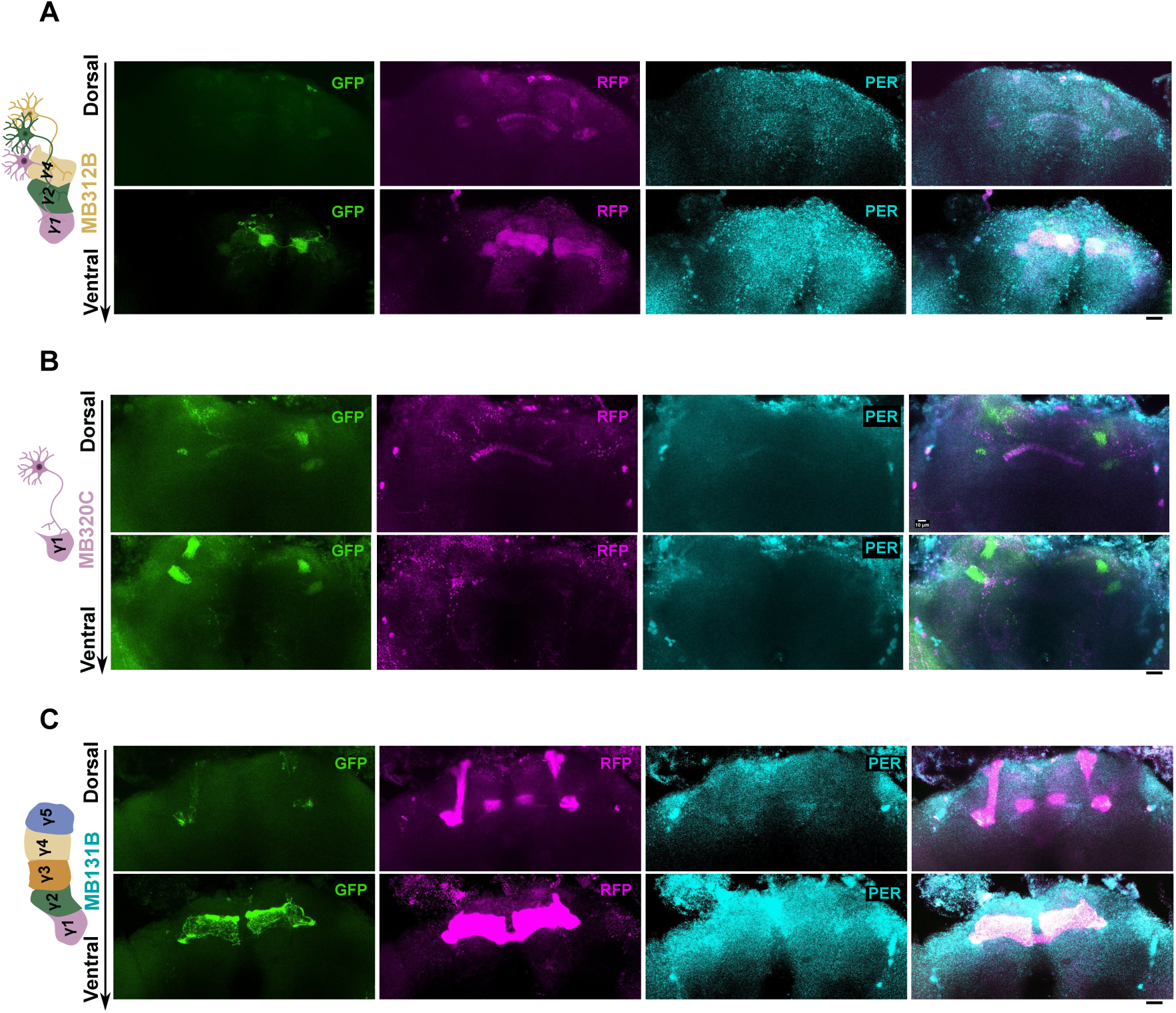
*Retro*-Tango tracing revealed no connectivity between clock neurons and PAM-γ1, -γ2, -γ4, PPL1, or γ-KCs. **A-C.** Representative images of *retro*-Tango-labeled neurons. The postsynaptic DA neuron subpopulations expressing GFP are shown in green, and presynaptic partners expressing mtdTomato are shown in magenta. Brains were co-stained with anti-PER antibodies to identify circadian clock neurons (cyan). No colocalization between the presynaptic partner and PER signal was observed. Scale bar, 20um. (**A**) *retro*-Tango conducted with the MB312B split GAL4-driver, targeting PAM-γ1, -γ2 and -γ4 neurons. (**B**) *retro*-Tango tracing with MB320C expressing in PPL1-γ1 peduncle neurons. (**C**) *retro*-Tango with the γ-KC driver MB131B.

**Supplementary Figure 5.**
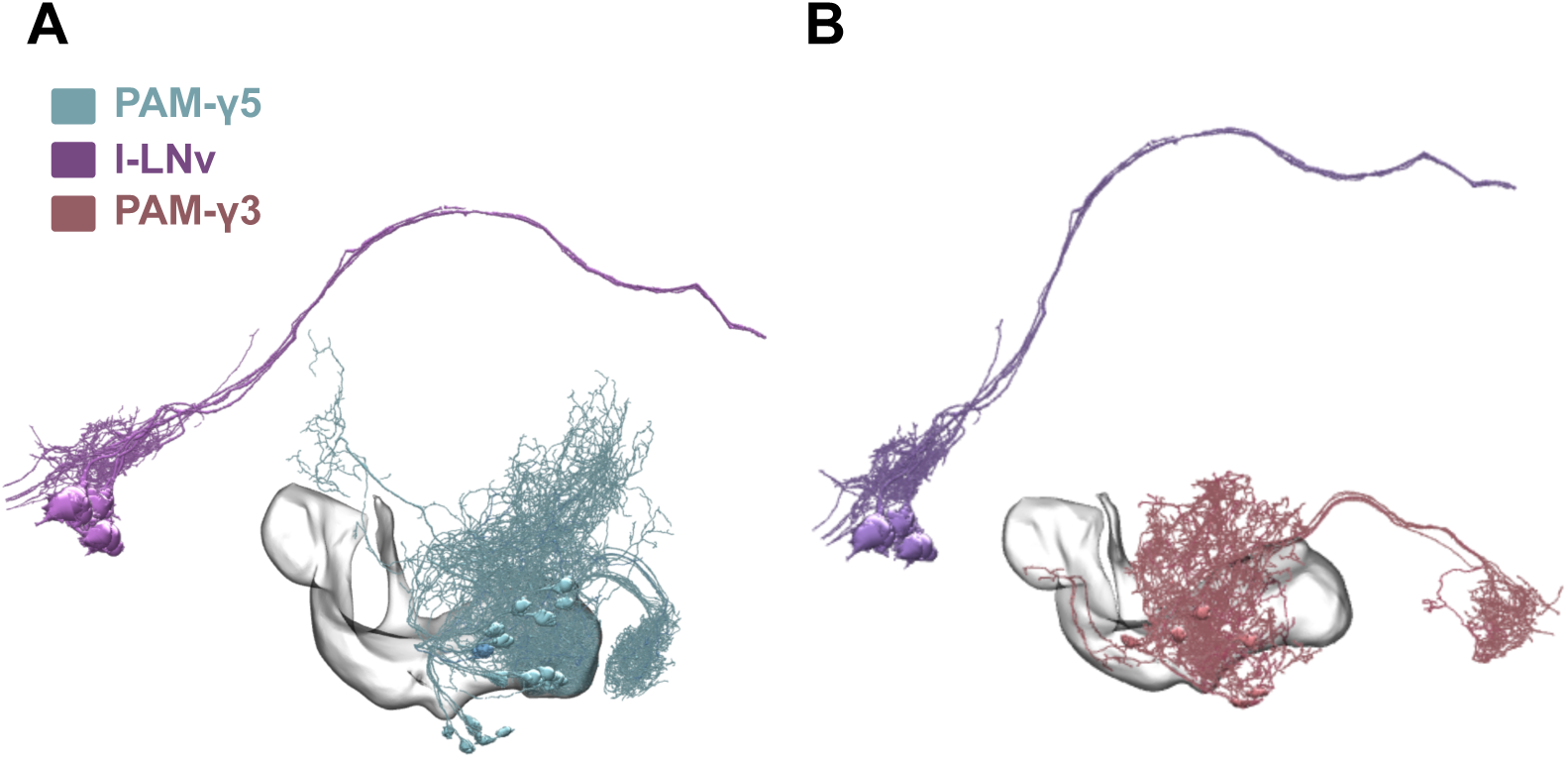
L-LNv clock neurons do not make physical contact with PAM-γ3 and -γ5 neurons. **A and B.** The hemibrain connectome data visualized using the NeuPrint tool. (**A**) Skeleton representation of the l-LNvs and PAM-γ3. (**B**) The l-LNvs and PAM-γ5 neurons.

### Supplementary Tables

**Supplementary Table 1.**
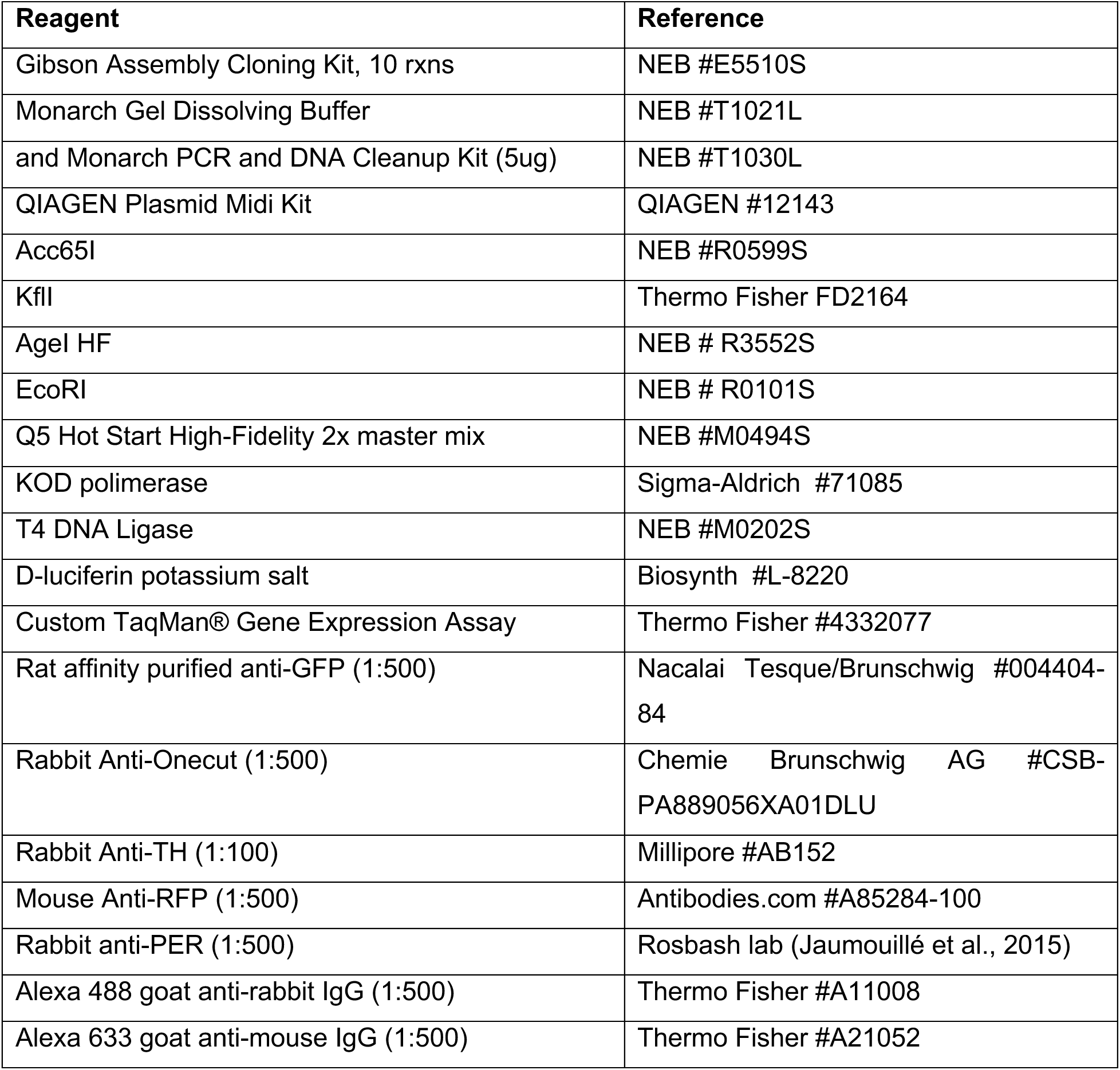
Description and references of the reagents and antibodies used in this study.

**Supplementary Table 2.**
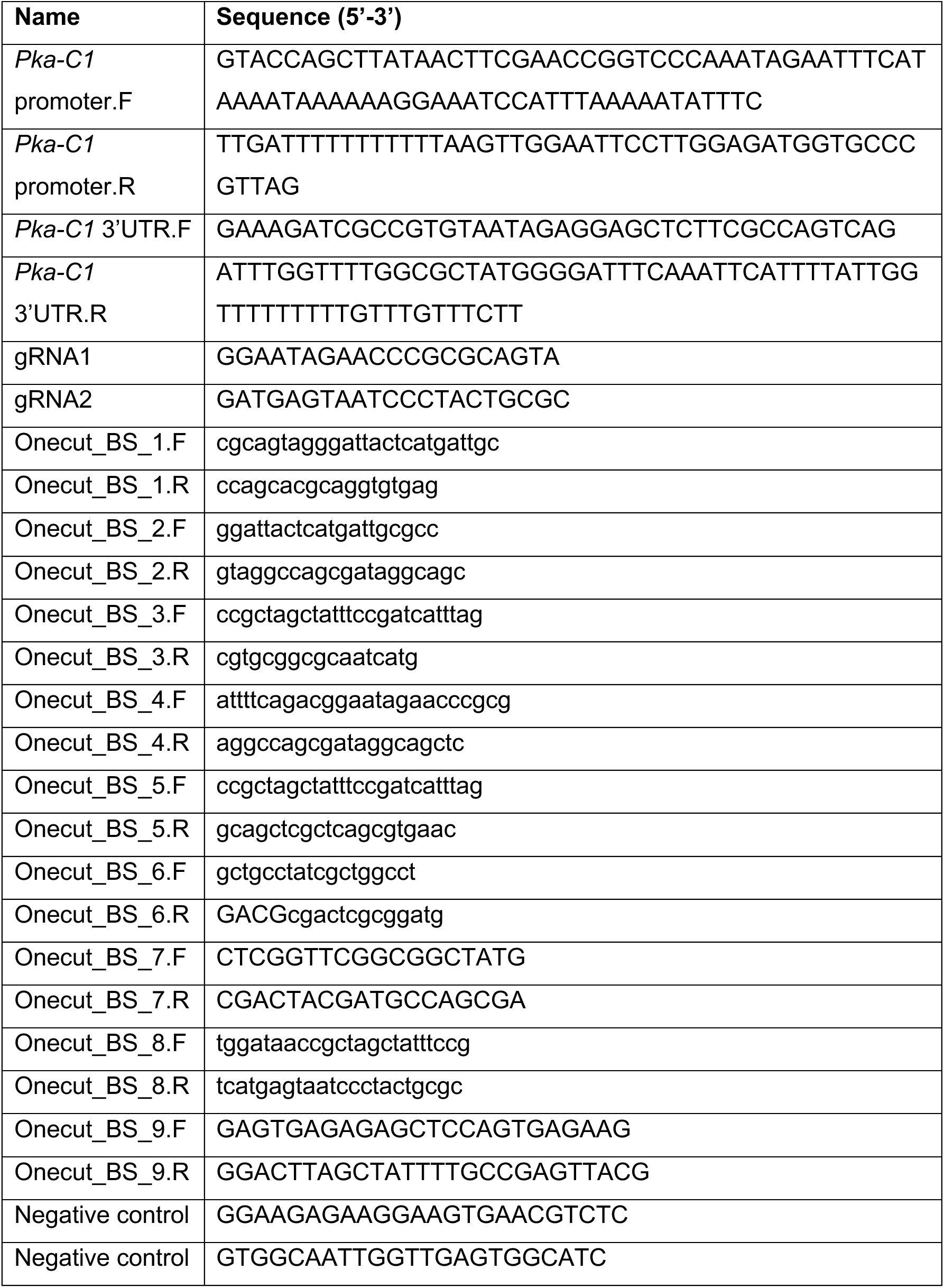
Primers and gRNAs used in this study.

**Supplementary Table 3.**
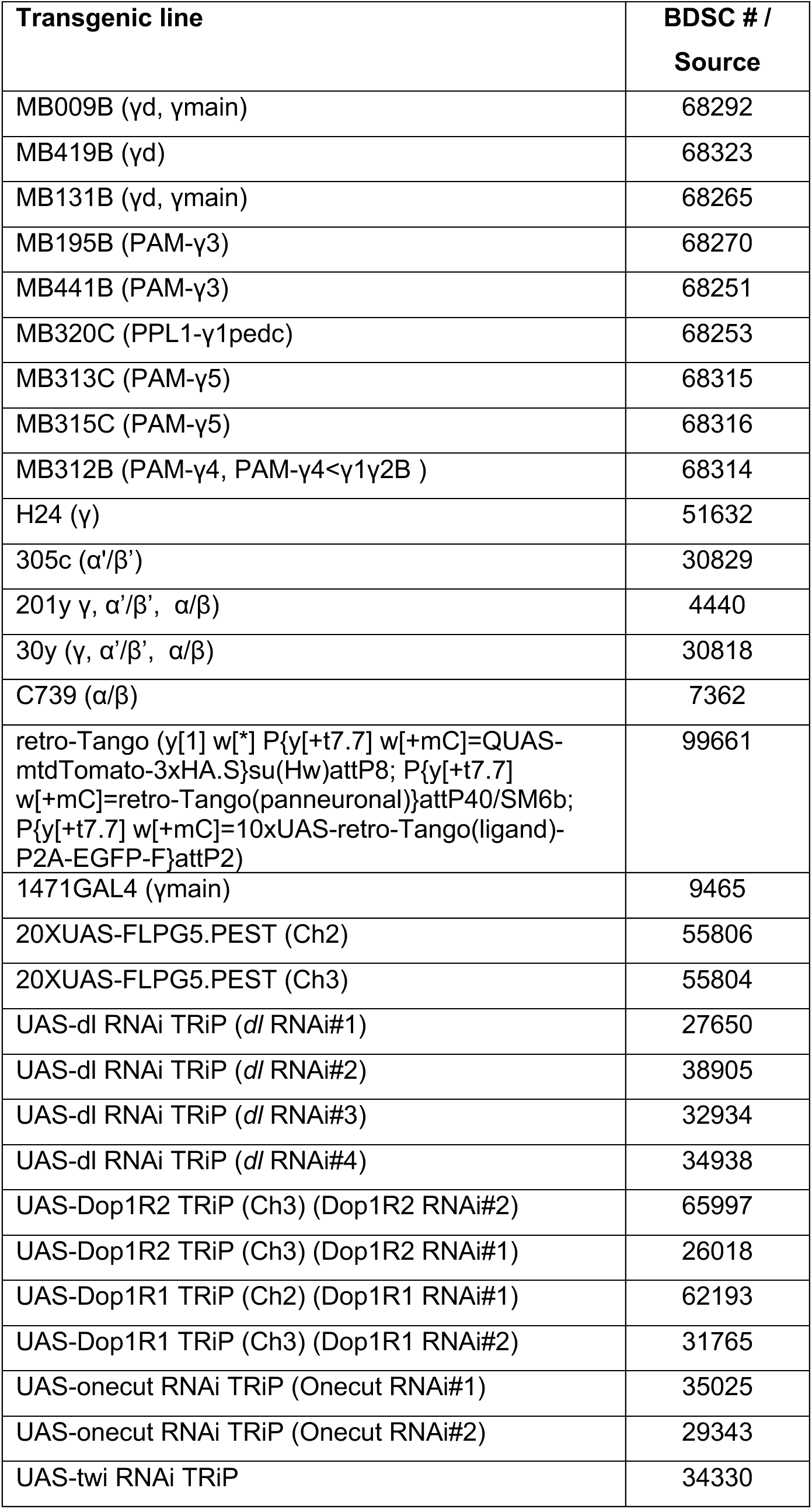
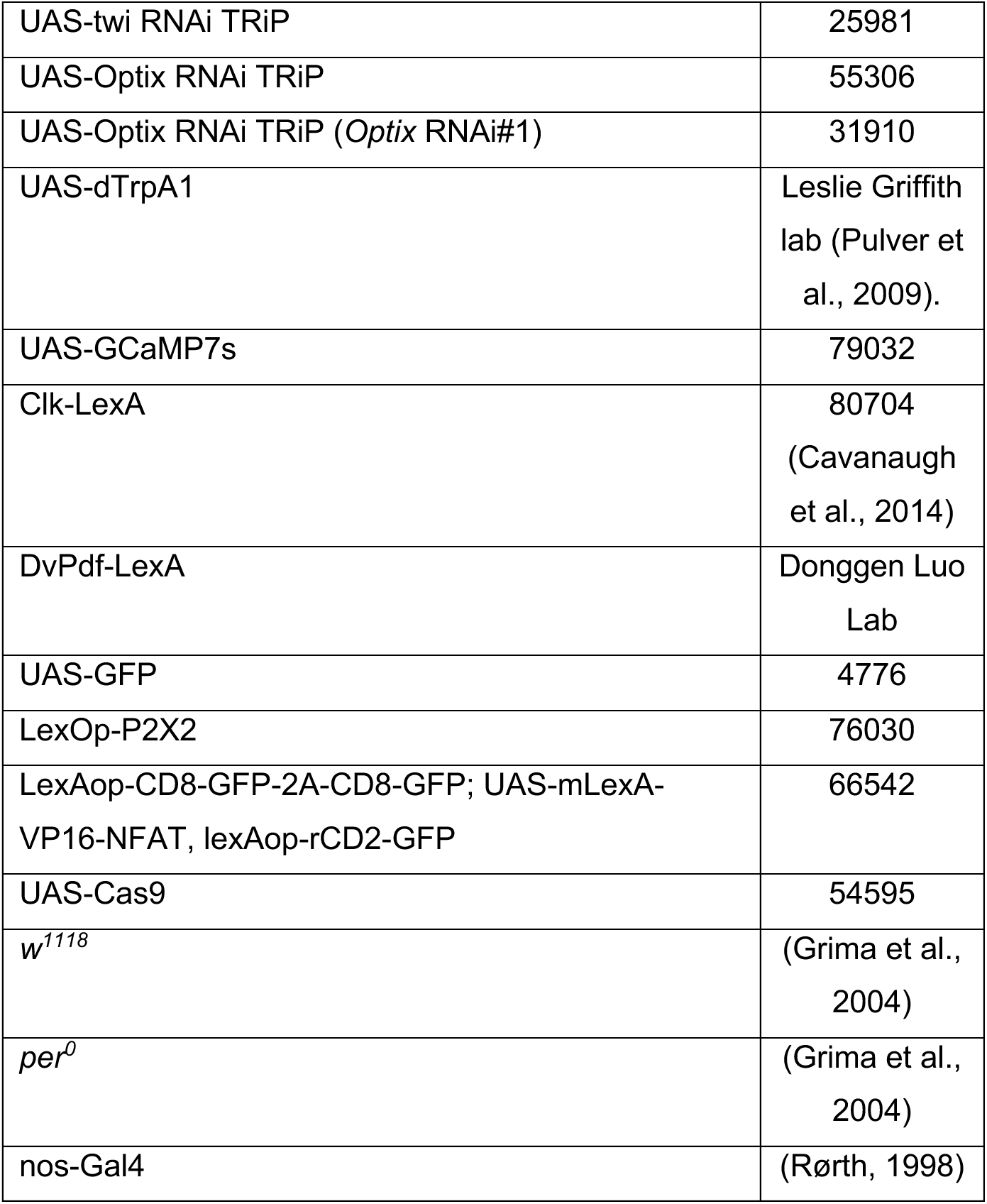
*Drosophila melanogaster* strains used in this study.

## REFERENCES

Abbott, S. M., & Zee, P. C. (2019). Circadian Rhythms: Implications for Health and Disease. Neurologic Clinics, 37(3), 601–613. 10.1016/J.NCL.2019.04.004

Allada, R., & Chung, B. Y. (2009). Circadian organization of behavior and physiology in Drosophila. Annual Review of Physiology, 72, 605–624. 10.1146/annurev-physiol-021909-135815

Artiushin, G., & Sehgal, A. (2017). The Drosophila circuitry of sleep–wake regulation. Current Opinion in Neurobiology, 44, 243–250. 10.1016/j.conb.2017.03.004

Ashmore, L. J., Sathyanarayanan, S., Silvestre, D. W., Emerson, M. M., Schotland, P., & Sehgal, A. (2003). Novel Insights into the Regulation of the Timeless Protein. The Journal of Neuroscience, 23(21), 7810–7819. 10.1523/JNEUROSCI.23-21-07810.2003

Aso, Y., Grübel, K., Busch, S., Friedrich, A. B., Siwanowicz, I., & Tanimoto, H. (2009). The mushroom body of adult Drosophila characterized by GAL4 drivers. Journal of Neurogenetics, 23(1–2), 156–172. 10.1080/01677060802471718

Aso, Y., Hattori, D., Yu, Y., Johnston, R. M., Iyer, N. A., Ngo, T.-T., Dionne, H., Abbott, L., Axel, R., Tanimoto, H., & Rubin, G. M. (2014). The neuronal architecture of the mushroom body provides a logic for associative learning. eLife, 3, e04577. 10.7554/eLife.04577

Aso, Y., Sitaraman, D., Ichinose, T., Kaun, K. R., Vogt, K., Belliart-Guérin, G., Plaçais, P.-Y., Robie, A. A., Yamagata, N., Schnaitmann, C., Rowell, W. J., Johnston, R. M., Ngo, T.-T. B., Chen, N., Korff, W., Nitabach, M. N., Heberlein, U., Preat, T., Branson, K. M., … Rubin, G. M. (2014). Mushroom body output neurons encode valence and guide memory-based action selection in Drosophila. eLife, 3, e04580. 10.7554/eLife.04580

Barber, A. F., Erion, R., Holmes, T. C., & Sehgal, A. (2016). Circadian and feeding cues integrate to drive rhythms of physiology in Drosophila insulin-producing cells. Genes and Development, 30(23), 2596–2606. 10.1101/gad.288258.116

Barber, A. F., Fong, S. Y., Kolesnik, A., Fetchko, M., & Sehgal, A. (2021). Drosophila clock cells use multiple mechanisms to transmit time-of-day signals in the brain. Proceedings of the National Academy of Sciences of the United States of America, 118(10), e2019826118. 10.1073/PNAS.2019826118/-/DCSUPPLEMENTAL

Bargiello, T. A., & Young, M. W. (1984). Molecular genetics of a biological clock in Drosophila. Proceedings of the National Academy of Sciences of the United States of America, *81*(7 I), 2142–2146. 10.1073/PNAS.81.7.2142

Campbell, S. S., & Tobler, I. (1984). Animal sleep: A review of sleep duration across phylogeny. Neuroscience and Biobehavioral Reviews, 8(3), 269–300. 10.1016/0149-7634(84)90054-X

Cavanaugh, D. J., Geratowski, J. D., Wooltorton, J. R. A., Spaethling, J. M., Hector, C. E., Zheng, X., Johnson, E. C., Eberwine, J. H., & Sehgal, A. (2014). Identification of a circadian output circuit for rest:activity rhythms in Drosophila. Cell, 157(3), 689–701. 10.1016/j.cell.2014.02.024

Chatterjee, A., Tanoue, S., Houl, J. H., & Hardin, P. E. (2010). Regulation of gustatory physiology and appetitive behavior by the circadian clock in Drosophila melanogaster. Current Biology : CB, 20(4), 300–309. 10.1016/j.cub.2009.12.055

Chung, B. Y., Ro, J., Hutter, S. A., Miller, K. M., Guduguntla, L. S., Kondo, S., & Pletcher, S. D. (2017). Drosophila Neuropeptide F Signaling Independently Regulates Feeding and Sleep-Wake Behavior. Cell Reports, 19(12), 2441–2450. 10.1016/j.celrep.2017.05.085

Davie, K., Janssens, J., Koldere, D., De Waegeneer, M., Pech, U., Kreft, Ł., Aibar, S., Makhzami, S., Christiaens, V., Bravo González-Blas, C., Poovathingal, S., Hulselmans, G., Spanier, K. I., Moerman, T., Vanspauwen, B., Geurs, S., Voet, T., Lammertyn, J., Thienpont, B., … Aerts, S. (2018). A Single-Cell Transcriptome Atlas of the Aging Drosophila Brain. Cell, 174(4), 982–998.e20. 10.1016/j.cell.2018.05.057

Depetris-Chauvin, A., Berni, J., Aranovich, E. J., Muraro, N. I., Beckwith, E. J., & Ceriani, M. F. (2011). Adult-Specific Electrical Silencing of Pacemaker Neurons Uncouples Molecular Clock from Circadian Outputs. Current Biology, 21(21), 1783–1793. 10.1016/j.cub.2011.09.027

Driscoll, M., Buchert, S. N., Coleman, V., McLaughlin, M., Nguyen, A., & Sitaraman, D. (2021). Compartment specific regulation of sleep by mushroom body requires GABA and dopaminergic signaling. Scientific Reports, 11(1), 1–18. 10.1038/s41598-021-99531-2

Duret, L. C., & Nagoshi, E. (2025). The intertwined relationship between circadian dysfunction and Parkinson’s disease. Trends in Neurosciences, 48(1), 62–76. 10.1016/j.tins.2024.10.006

Fropf, R., Zhou, H., & Yin, J. C. P. (2018). The clock gene period differentially regulates sleep and memory in Drosophila. Neurobiology of Learning and Memory, 153, 2–12. 10.1016/J.NLM.2018.02.016

Garczynski, S. F., Brown, M. R., Shen, P., Murray, T. F., & Crim, J. W. (2002). Characterization of a functional neuropeptide F receptor from Drosophila melanogaster. Peptides, 23(4), 773–780. 10.1016/s0196-9781(01)00647-7

Glossop, N. R. J., Houl, J. H., Zheng, H., Ng, F. S., Dudek, S. M., & Hardin, P. E. (2003). VRILLE Feeds Back to Control Circadian Transcription of Clock in the Drosophila Circadian Oscillator. Neuron, 37(2), 249–261. 10.1016/S0896-6273(03)00002-3

Grima, B., Chélot, E., Xia, R., & Rouyer, F. (2004). Morning and evening peaks of activity rely on different clock neurons of the Drosophila brain. Nature, 431(7010), 869–873. 10.1038/nature02935

Handler, A., Graham, T. G. W., Cohn, R., Morantte, I., Siliciano, A. F., Zeng, J., Li, Y., & Ruta, V. (2019). Distinct dopamine receptor pathways underlie the temporal sensitivity of associative learning. Cell, 178(1), 60–75.e19. 10.1016/j.cell.2019.05.040

Hattori, D., Aso, Y., Swartz, K. J., Rubin, G. M., Abbott, L. F., & Axel, R. (2017). Representations of Novelty and Familiarity in a Mushroom Body Compartment. Cell, 169(5), 956–969.e17. 10.1016/j.cell.2017.04.028

Jabbur, M. L., Dani, C., Spoelstra, K., Dodd, A. N., & Johnson, C. H. (2024). Evaluating the Adaptive Fitness of Circadian Clocks and their Evolution. Journal of Biological Rhythms, 39(2), 115–134. 10.1177/07487304231219206

Jafari, S., Alkhori, L., Schleiffer, A., Brochtrup, A., Hummel, T., & Alenius, M. (2012). Combinatorial Activation and Repression by Seven Transcription Factors Specify Drosophila Odorant Receptor Expression. PLoS Biology, 10(3), e1001280. 10.1371/journal.pbio.1001280

Jauregui-Lozano, J., Hall, H., Stanhope, S. C., Bakhle, K., Marlin, M. M., & Weake, V. M. (2022). The Clock:Cycle complex is a major transcriptional regulator of Drosophila photoreceptors that protects the eye from retinal degeneration and oxidative stress. PLoS Genetics, 18(1), e1010021. 10.1371/journal.pgen.1010021

Joiner, W. J., Crocker, A., White, B. H., & Sehgal, A. (2006). Sleep in Drosophila is regulated by adult mushroom bodies. Nature, 441(7094), 757–760. 10.1038/nature04811

Kayser, M. S., Yue, Z., & Sehgal, A. (2014). A critical period of sleep for development of courtship circuitry and behavior in Drosophila. Science, 344(6181), 269–274. 10.1126/science.1250553

Keder, A., Tardieu, C., Malong, L., Filia, A., Kashkenbayeva, A., Newton, F., Georgiades, M., Gale, J. E., Lovett, M., Jarman, A. P., & Albert, J. T. (2020). Homeostatic maintenance and age-related functional decline in the Drosophila ear. Scientific Reports, 10(1), 7431. 10.1038/s41598-020-64498-z

Kim, Y.-C., Lee, H.-G., & Han, K.-A. (2007). D1 Dopamine Receptor dDA1 Is Required in the Mushroom Body Neurons for Aversive and Appetitive Learning in Drosophila. The Journal of Neuroscience, 27(29), 7640–7647. 10.1523/JNEUROSCI.1167-07.2007

Kume, K., Kume, S., Park, S. K., Hirsh, J., & Jackson, F. R. (2005). Dopamine is a regulator of arousal in the fruit fly. The Journal of Neuroscience: The Official Journal of the Society for Neuroscience, 25(32), 7377–7384. 10.1523/JNEUROSCI.2048-05.2005

Lago Solis, B., Koch, R., & Nagoshi, E. (2025). Circadian clock-independent ultradian rhythms in lipid metabolism in the *Drosophila* fat body. Journal of Biological Chemistry, 301(6), 110245. 10.1016/j.jbc.2025.110245

Li, F., Lindsey, J. W., Marin, E. C., Otto, N., Dreher, M., Dempsey, G., Stark, I., Bates, A. S., Pleijzier, M. W., Schlegel, P., Nern, A., Takemura, S.-Y., Eckstein, N., Yang, T., Francis, A., Braun, A., Parekh, R., Costa, M., Scheffer, L. K., … Rubin, G. M. (2020). The connectome of the adult Drosophila mushroom body provides insights into function. eLife, 9, e62576. 10.7554/eLife.62576

Li, H., Janssens, J., De Waegeneer, M., Kolluru, S. S., Davie, K., Gardeux, V., Saelens, W., David, F. P. A., Brbić, M., Spanier, K., Leskovec, J., McLaughlin, C. N., Xie, Q., Jones, R. C., Brueckner, K., Shim, J., Tattikota, S. G., Schnorrer, F., Rust, K., … Zinzen, R. P. (2022). Fly Cell Atlas: A single-nucleus transcriptomic atlas of the adult fruit fly. *Science (New York*, N.Y.), 375(6584), eabk2432. 10.1126/science.abk2432

Liang, X., Holy, T. E., & Taghert, P. H. (2016). Synchronous Drosophila circadian pacemakers display non-synchronous Ca2+ rhythms in vivo. *Science (New York*, N.Y.), 351(6276), 976–981. 10.1126/science.aad3997

Liu, W., Ganguly, A., Huang, J., Wang, Y., Ni, J. D., Gurav, A. S., Aguilar, M. A., & Montell, C. (n.d.). Neuropeptide F regulates courtship in Drosophila through a male-specific neuronal circuit. eLife, 8, e49574. 10.7554/eLife.49574

Machado Almeida, P., Lago Solis, B., Stickley, L., Feidler, A., & Nagoshi, E. (2021). Neurofibromin 1 in mushroom body neurons mediates circadian wake drive through activating cAMP-PKA signaling. Nature Communications, 12(1), 5758. 10.1038/s41467-021-26031-2

Majcin Dorcikova, M., Duret, L. C., Pottié, E., & Nagoshi, E. (2023). Circadian clock disruption promotes the degeneration of dopaminergic neurons in male Drosophila. Nature Communications, 14(1). 10.1038/s41467-023-41540-y

Mao, Z., & Davis, R. L. (2009). Eight Different Types of Dopaminergic Neurons Innervate the Drosophila Mushroom Body Neuropil: Anatomical and Physiological Heterogeneity. Frontiers in Neural Circuits, 3, 5. 10.3389/neuro.04.005.2009

Martin-Burgos, B., Wang, W., William, I., Tir, S., Mohammad, I., Javed, R., Smith, S., Cui, Y., Arzavala, J., Mora, D., Smith, C. B., van der Vinne, V., Molyneux, P. C., Miller, S. C., Weaver, D. R., Leise, T. L., & Harrington, M. E. (2022). Methods for Detecting PER2:LUCIFERASE Bioluminescence Rhythms in Freely Moving Mice. Journal of Biological Rhythms, 37(1), 78–93. 10.1177/07487304211062829

Masuyama, K., Zhang, Y., Rao, Y., & Wang, J. W. (2012). Mapping neural circuits with activity-dependent nuclear import of a transcription factor. Journal of Neurogenetics, 26(1), 89–102. 10.3109/01677063.2011.642910

Modi, M. N., Shuai, Y., & Turner, G. C. (2020). The Drosophila Mushroom Body: From Architecture to Algorithm in a Learning Circuit. Annual Review of Neuroscience, 43, 465–484. 10.1146/annurev-neuro-080317-0621333

Muraro, N. I., Pírez, N., & Ceriani, M. F. (2013). The circadian system: Plasticity at many levels. Neuroscience, 247, 280–293. 10.1016/j.neuroscience.2013.05.036

Nguyen, D. N. T., Rohrbaugh, M., & Lai, Z.-C. (2000). The *Drosophila* homolog of Onecut homeodomain proteins is a neural-specific transcriptional activator with a potential role in regulating neural differentiation. Mechanisms of Development, 97(1), 57–72. 10.1016/S0925-4773(00)00431-7

Peschel, N., & Helfrich-Förster, C. (2011). Setting the clock—By nature: Circadian rhythm in the fruitfly Drosophila melanogaster. FEBS Letters, 585(10), 1435–1442. 10.1016/j.febslet.2011.02.028

Qin, H., Cressy, M., Li, W., Coravos, J. S., Izzi, S. A., & Dubnau, J. (2012). Gamma neurons mediate dopaminergic input during aversive olfactory memory formation in Drosophila. Current Biology: CB, 22(7), 608–614. 10.1016/j.cub.2012.02.014

Schindelin, J., Arganda-Carrera, I., Frise, E., Verena, K., Mark, L., Tobias, P., Stephan, P., Curtis, R., Stephan, S., Benjamin, S., Jean-Yves, T., Daniel, J. W., Volker, H., Kevin, E., Pavel, T., & Albert, C. (2009). Fiji—An Open platform for biological image analysis. Nature Methods, 9(7). 10.1038/nmeth.2019.Fiji

Schlegel, P., Yin, Y., Bates, A. S., Dorkenwald, S., Eichler, K., Brooks, P., Han, D. S., Gkantia, M., dos Santos, M., Munnelly, E. J., Badalamente, G., Serratosa Capdevila, L., Sane, V. A., Fragniere, A. M. C., Kiassat, L., Pleijzier, M. W., Stürner, T., Tamimi, I. F. M., Dunne, C. R., … Jefferis, G. S. X. E. (2024). Whole-brain annotation and multi-connectome cell typing of Drosophila. Nature, 634(8032), 139–152. 10.1038/s41586-024-07686-5

Schubert, F. K., Hagedorn, N., Yoshii, T., Helfrich-Förster, C., & Rieger, D. (2018). Neuroanatomical details of the lateral neurons of Drosophila melanogaster support their functional role in the circadian system. The Journal of Comparative Neurology, 526(7), 1209–1231. 10.1002/cne.24406

Schwaerzel, M., Monastirioti, M., Scholz, H., Friggi-Grelin, F., Birman, S., & Heisenberg, M. (2003). Dopamine and Octopamine Differentiate between Aversive and Appetitive Olfactory Memories in Drosophila. The Journal of Neuroscience, 23(33), 10495–10502. 10.1523/JNEUROSCI.23-33-10495.2003

Shang, Y., Donelson, N. C., Vecsey, C. G., Guo, F., Rosbash, M., & Griffith, L. C. (2013). Short neuropeptide F is a sleep-promoting inhibitory modulator. Neuron, 80(1), 171–183. 10.1016/j.neuron.2013.07.029

Shaw, P. J., Cirelli, C., Greenspan, R. J., & Tononi, G. (2000). Correlates of sleep and waking in Drosophila melanogaster. *Science (New York*, N.Y.), 287(5459), 1834–1837. 10.1126/science.287.5459.1834

Siju, K. P., De Backer, J.-F., & Grunwald Kadow, I. C. (2021). Dopamine modulation of sensory processing and adaptive behavior in flies. Cell and Tissue Research, 383(1), 207–225. 10.1007/s00441-020-03371-x

Sorkaç, A., Moșneanu, R. A., Crown, A. M., Savaş, D., Okoro, A. M., Memiş, E., Talay, M., & Barnea, G. (2023). Retro-Tango enables versatile retrograde circuit tracing in Drosophila. eLife, 12, 1–18. 10.7554/eLife.85041

Streeper, R. S., Hornbuckle, L. A., Svitek, C. A., Goldman, J. K., Oeser, J. K., & O’Brien, R. M. (2001). Protein kinase A phosphorylates hepatocyte nuclear factor-6 and stimulates glucose-6-phosphatase catalytic subunit gene transcription. The Journal of Biological Chemistry, 276(22), 19111–19118. 10.1074/jbc.M101442200

Sun, J., Xu, A. Q., Giraud, J., Poppinga, H., Riemensperger, T., Fiala, A., & Birman, S. (2018). Neural Control of Startle-Induced Locomotion by the Mushroom Bodies and Associated Neurons in Drosophila. Frontiers in Systems Neuroscience, 12, 6. 10.3389/fnsys.2018.00006

Swinderen, B. van. (2009). Fly Memory: A Mushroom Body Story in Parts. Current Biology, 19(18), R855–R857. 10.1016/j.cub.2009.07.064

Vassalli, Q. A., Colantuono, C., Nittoli, V., Ferraioli, A., Fasano, G., Berruto, F., Chiusano, M. L., Kelsh, R. N., Sordino, P., & Locascio, A. (2021). Onecut Regulates Core Components of the Molecular Machinery for Neurotransmission in Photoreceptor Differentiation. Frontiers in Cell and Developmental Biology, 9, 602450. 10.3389/fcell.2021.602450

Woelfle, M. A., Ouyang, Y., Phanvijhitsiri, K., & Johnson, C. H. (2004). The adaptive value of circadian clocks: An experimental assessment in cyanobacteria. Current Biology, 14(16), 1481–1486. 10.1016/J.CUB.2004.08.023

Wu, F., Li, R., Umino, Y., Kaczynski, T. J., Sapkota, D., Li, S., Xiang, M., Fliesler, S. J., Sherry, D. M., Gannon, M., Solessio, E., & Mu, X. (2013). Onecut1 Is Essential for Horizontal Cell Genesis and Retinal Integrity. The Journal of Neuroscience, 33(32), 13053–13065. 10.1523/JNEUROSCI.0116-13.2013

Xue, Y., & Zhang, Y. (2018). Emerging roles for microRNA in the regulation of Drosophila circadian clock. BMC Neuroscience, 19(1), 1. 10.1186/s12868-018-0401-8

Yang, M., Lee, J.-E., Padgett, R. W., & Edery, I. (2008). Circadian regulation of a limited set of conserved microRNAs in Drosophila. BMC Genomics, 9, 83. 10.1186/1471-2164-9-83

Yao, Z., Macara, A. M., Lelito, K. R., Minosyan, T. Y., & Shafer, O. T. (2012). Analysis of functional neuronal connectivity in the Drosophila brain. Journal of Neurophysiology, 108(2), 684–696. 10.1152/jn.00110.2012

Zars, T., Fischer, M., Schulz, R., & Heisenberg, M. (2000). Localization of a short-term memory in Drosophila. *Science (New York*, N.Y.), 288(5466), 672–675. 10.1126/science.288.5466.672

Zhou, J., Yu, W., & Hardin, P. E. (2015). ChIPping Away at the Drosophila Clock. In A. Sehgal (Ed.), Methods in Enzymology (Vol. 551, pp. 323–347). Academic Press. 10.1016/bs.mie.2014.10.019

Zordan, M. A., & Sandrelli, F. (2015). Circadian clock dysfunction and psychiatric disease: Could fruit flies have a say? Frontiers in Neurology, 6(MAR). 10.3389/fneur.2015.00080

## References

Jaumouillé, E., Machado Almeida, P., Stähli, P., Koch, R., & Nagoshi, E. (2015). Transcriptional regulation via nuclear receptor crosstalk required for the drosophila circadian clock. Current Biology, 25(11), 1502–1508. 10.1016/j.cub.2015.04.017

Pulver, S. R., Pashkovski, S. L., Hornstein, N. J., Garrity, P. A., & Griffith, L. C. (2009). Temporal dynamics of neuronal activation by Channelrhodopsin-2 and TRPA1 determine behavioral output in Drosophila larvae. Journal of Neurophysiology, 101(6), 3075–3088. 10.1152/jn.00071.2009

Rørth, P. (1998). Gal4 in the Drosophila female germline. Mechanisms of Development, 78(1– 2), 113–118. 10.1016/s0925-4773(98)00157-9

